# Global analysis of the RNA-RNA interactome in *Acinetobacter baumannii* AB5075 uncovers a small regulatory RNA repressing the virulence-related outer membrane protein CarO

**DOI:** 10.1101/2023.12.04.569942

**Authors:** Fergal J. Hamrock, Daniel Ryan, Ali Shaibah, Anna S. Ershova, Aalap Mogre, Maha M. Sulimani, Sarah Reichardt, Karsten Hokamp, Alexander J. Westermann, Carsten Kröger

**Affiliations:** Department of Microbiology, School of Genetics & Microbiology, Moyne Institute of Preventive Medicine, Trinity College Dublin, Dublin 2, Ireland; Helmholtz Institute for RNA-based Infection Research (HIRI), Helmholtz Centre for Infection Research (HZI), Würzburg, Germany; Department of Genetics, School of Genetics & Microbiology, Smurfit Institute of Genetics, Trinity College Dublin, Dublin 2, Ireland; Institute of Molecular Infection Biology (IMIB), University of Würzburg, Würzburg, Germany; Department of Microbiology, Biocentre, University of Würzburg, Würzburg, Germany

**Keywords:** Hi-GRIL-seq, RNA-RNA interactome, *A. baumannii*, small RNA, Aar, CarO, translational repression

## Abstract

*Acinetobacter baumannii* is an opportunistic Gram-negative pathogen that infects critically ill patients. The emergence of antimicrobial resistant *A. baumannii* has exacerbated the need to functionally characterise environmental adaptation, antibiotic resistance and pathogenicity of this organism and their genetic regulators to inform intervention strategies. Critical to rapid adaptation to changing environments in bacteria are small regulatory RNAs (sRNAs), however, the role that sRNAs play in the biology of *A. baumannii* is poorly understood. To assess the regulatory function of sRNAs and to uncover their RNA interaction partners in *A. baumannii*, we employed an RNA proximity ligation and sequencing method (Hi-GRIL-seq) in three different environmental conditions. We found that 40 sRNA candidates were ligated to sRNA-RNA chimeric sequencing reads, suggesting that sRNA-mediated gene regulation is pervasive in *A. baumannii* and that sRNAs act as direct regulators of mRNA molecules through antisense base-pairing. In-depth characterisation uncovered the sRNA Aar to be a post-transcriptional regulator of four mRNA targets including that of the outer membrane protein CarO and the siderophore receptor BfnH. We show that Aar initiates base-pairing with these mRNA molecules using a conserved seed region of nine nucleotides, sequestering the ribosome binding sites and inhibiting translation. Aar is differentially expressed in response to multiple stress stimuli suggesting a role in fine-tuning translation of the Aar-target molecules in *A. baumannii* under hostile conditions. Together, our study provides mechanistic insights into sRNA-mediated gene expression control in *A. baumannii* and represents a valuable resource for future RNA-centric research endeavours in this ESKAPE pathogen.

## INTRODUCTION

The opportunistic pathogen *A. baumannii* is considered a serious threat to human and animal health because of high-level antimicrobial resistance (AMR) (Tacconelli *et al*., 2018; Whiteway *et al*., 2021; Cain and Hamidian, 2023). The World Health Organization has declared *A. baumannii* as a priority organism that demands investment into research and development of novel antibiotics (Tacconelli *et al*., 2018). Diseases caused by this pathogen—namely ventilator-associated pneumonia, urinary tract and bloodstream infections—are commonly associated with intrusive medical devices and surgery (Dijkshoorn *et al*., 2007; Harding *et al*., 2018). To rapidly adapt to changing environmental conditions, *A. baumannii* alters the expression of critical stress response genes (Kröger *et al*., 2016; Wood *et al*., 2018). The role that protein-based transcriptional regulators exert in response to environmental stressors has been the subject of various studies, however, the role that post-transcriptional regulation plays in controlling *A. baumannii* gene expression is largely unknown, yet would offer much needed insight into the biology of this pathogen (Weiss *et al*., 2016; Kröger *et al*., 2016; Kröger *et al*., 2018; Parmeciano Di Noto *et al*., 2019).

Post-transcriptional regulation in bacteria is often mediated by the action of non-coding, small regulatory RNAs (sRNAs) that exert important physiological roles such as controlling AMR, virulence, metabolism, envelope stress and iron starvation responses (Papenfort and Vogel, 2014; Sharma C. M. and Svensson S., 2016; Dersch *et al*., 2017; Chareyre and Mandin, 2018; Westermann, 2018; Papenfort and Melamed, 2023). Initial contacts between sRNAs and their cognate mRNA targets are mediated by conserved contiguous “seed” nucleotides in the sRNA molecule that bind to complementary regions of the mRNA targets (Gottesman and Storz, 2011; Gorski *et al*., 2017). Most known sRNAs inhibit translation initiation of their mRNA targets by occluding the small ribosomal subunit from binding to the Shine Dalgarno (SD) sequence or preventing ribosome progression through the first five codons of the target mRNA, which may be accompanied by the destabilisation of the untranslated target transcript (Storz *et al*., 2011; Wagner and Romby, 2015; Papenfort and Melamed, 2023).

Our knowledge of the role that sRNAs play in the regulatory mechanisms of *Acinetobacter* spp. is still in its infancy. The first putative regulatory sRNA Aar (<u>A</u>mino <u>a</u>cid regulator) was identified in non-pathogenic *Acinetobacter baylyi* and is conserved in *A. baumannii* (Schilling *et al*., 2010). A global bioinformatic analysis additionally predicted 31 candidate sRNAs in *A. baumannii*, three of which were validated by Northern blotting (Sharma *et al*., 2014). More recently, transcriptomic analyses increased the number of candidate sRNAs in *A. baumannii* AB5075 to 88 (six validated by Northern blotting) and in *A. baumannii* ATCC17978 to 110 (seven validated by Northern blotting) (Weiss *et al*., 2016; Kröger *et al*., 2018). Only seven *A. baumannii* ATCC17978 sRNAs are conserved (between 70-75% sequence identity) across selected members of the *Pseudomonadales* (e.g., *Pseudomonas aeruginosa* and *Moraxella catarrhalis*) and none shares any sequence conservation with sRNAs in *Enterobactericeae* (Kröger *et al*., 2018). A few sRNAs in *A. baumannii* have been functionally characterised and were shown to be involved in the regulation of efflux pumps, biofilm formation and phenotypic switching between virulent and avirulent phase variants (Sharma *et al*., 2014; Álvarez-Fraga *et al*., 2017; Anderson *et al*., 2020; Pérez-Varela *et al*., 2022; Shenkutie *et al*., 2023). However, their direct targets and mechanism of action have not been dissected.

Typically, the activity of trans-encoded sRNAs in Gram-negative bacteria depends on RNA-binding proteins (RBPs) (Vogel and Luisi, 2011; Holmqvist and Vogel, 2018; Felden and Augagneur, 2021; Olejniczak *et al*., 2022; Liao and Smirnov, 2023; Papenfort and Melamed, 2023). For example, in *Enterobacteriaceae* the protein Hfq is frequently required for mediating sRNA-mRNA annealing (Vogel and Luisi, 2011; Santiago-Frangos and Woodson, 2018). In stark contrast, the role that RBPs play for sRNA-mediated control in *Acinetobacter* spp. is poorly understood. While an Hfq homologue is present *Acinetobacter* spp., the protein adopts an unusual structure compared to other bacterial species, with an enlarged, glycine rich C-terminal domain (Schilling and Gerischer, 2009). Ectopic expression of *A. baylyi* Hfq in an *hfq* deletion strain of *E. coli* restored growth and phenotype to wild-type levels, suggesting that *Acinetobacter* Hfq has RNA-RNA matchmaking capabilities (Schilling and Gerischer, 2009). Indeed, deletion of *hfq* in *A. baumannii* causes pleiotropic phenotypes with reduced growth, broad environmental sensitivity and attenuated virulence (Kuo *et al*., 2017; Sharma *et al*., 2018). Further, purified Hfq from *A. baumannii* was demonstrated to bind *A. baumannii* sRNA AbsR25 (sRNA65) and *E. coli* sRNAs MicA and DsrA *in vitro* (Sharma *et al*., 2018) as well as facilitated regulation of the *sodB* mRNA from *A. baumannii* by the known enterobacterial sRNA RyhB (Sharma *et al*., 2018). Despite these observations, an endogenous role of *A. baumannii* Hfq as a global RNA-RNA matchmaker in its native host has not been proven.

In the present study, we harnessed Hi-GRIL-seq—an *in vivo* RNA proximity ligation approach—to map the sRNA-mRNA interaction network of *A. baumannii* AB5075 in a global, yet RBP-independent manner. We found that of the 88 sRNA candidates annotated in *A. baumannii* AB5075 (Kröger *et al*., 2018), 40 were present in RNA-RNA chimeras, giving rise to 706 putative sRNA-mRNA interaction pairs. We selected the *A. baumannii* sRNA Aar as an illustrative example to highlight the value of this Hi-GRIL-seq dataset. We found that Aar was ligated to four likely mRNA targets, including the transcripts encoding the outer membrane protein (OMP) CarO and the TonB-dependent siderophore receptor BfnH. We confirmed these mRNA interactions using a combination of genetic and biochemical approaches. Additionally, we revealed that Aar employs a universal seed region of nine nucleotides to bind and regulate each of these four mRNA targets, providing us with the first mechanistic insight into sRNA-mediated post-transcriptional regulation in *A. baumannii*. Furthermore, we demonstrated that Aar expression is induced in different environmental conditions, suggesting that it may fine-tune gene expression in *Acinetobacter* species in response to cellular stress. We anticipate that our dataset will serve as a starting point to uncover many more biological roles for sRNA-mediated gene regulation in *A. baumannii*. To facilitate this, we have made our data easily accessible in an online browser at: http://bioinf.gen.tcd.ie/jbrowse2/Hi-GRIL-seq.

## MATERIAL AND METHODS

### Bacterial strains and growth conditions

*Acinetobacter baumannii* AB5075 and *Escherichia coli* TOP10 (Invitrogen) were maintained on lysogeny broth (Lennox (L-) agar plates (10 g/L tryptone, 5 g/L yeast extract, 5 g/L NaCl, 15 g/L agar)) and grown over night in lysogeny broth (Lennox (LB-), 10 g/L tryptone, 5 g/L yeast extract, 5 g/L NaCl) (Jacobs *et al*., 2014). Media were supplemented with the zeocin (*A. baumannii*: 250 μg/mL, *E. coli*: 10 μg/mL), tetracycline (12 μg/mL), ampicillin (150 μg/mL), apramycin (60 μg/mL) or chloramphenicol (25 μg/mL), sucrose (20% w/v) when necessary. To assess expression of Aar in response to changes in environmental conditions, *A. baumannii* AB5075 was first grown to late exponential phase (OD_600_ = 1, 220 rpm, 25 mL) at either 25°C or 37°C. These cultures were then “shocked” for 15 min prior to RNA isolation. The cultures grown at 25°C were heat-shocked by transferring flasks to 37°C. The samples grown at 37°C were either osmo-shocked with the addition of NaCl (0.3 M, final concentration), iron-deprived with 2,2′-dipyridyl (0.2 mM, final concentration) or cell envelope compromised with polymyxin B (1 µg/mL, final concentration). Negative controls where the cells were grown to late exponential phase at 25°C or 37°C before being returned to grow at their respective temperature for 15 min prior to RNA isolation were also included. The expression of Aar in these samples was detected using Northern blotting (see below). The relative expression of Aar in each sample was then quantified by measuring Aar signal intensity relative to the 5S rRNA loading control with ImageJ (Schneider *et al*., 2012).

### Generation of *A. baumannii* mutants

Deletion of *aar* (sRNA21) in *A. baumannii* AB5075 was accomplished by homologous recombination (Kröger *et al*., 2018; Godeux *et al*., 2020). The oligonucleotides used in this study are listed in **Supplementary Table 1**. Approximately 2,000 bp regions up- and downstream of *aar* were amplified using genomic DNA isolated from *A. baumannii* AB5075 as a template (primers 1 & 2 and 3 & 4). The *sacB*-*aacC4* genes were amplified from the pMHL2 plasmid (primers 5 & 6) (Godeux *et al*., 2020). A chimeric PCR product composed of the *sacB*-*aacC4* genes flanked by the upstream and downstream regions was constructed using overlap extension PCR (primers 1 & 4). Wild-type *A. baumannii* AB5075 were transformed with this chimeric PCR product using natural transformation (Godeux *et al*., 2018; Godeux *et al*., 2020). Colonies containing the insertion product were screened for apramycin resistance and sucrose sensitivity. The incorporation of the *sacB*-*aacC4* cassette was validated by colony PCR (primers 1 & 4). A second chimeric PCR product, composed solely of the ∼2,000 bp upstream and downstream regions, was created by overlap extension PCR (primers 1 & 7 and 4 & 8, respectively) and the *A. baumannii* Δ*aar::sacB_aacC4* insertion mutant strain was transformed with this PCR product to create a scarless *A. baumannii* AB5075 Δ*aar* mutant strain. Obtained colonies were screened for sucrose resistance and apramycin sensitivity. Loss of the *aar* gene was validated by PCR (using primers 9 & 10) and by whole-genome sequencing (MinION sequencer, Oxford Nanopore Technologies). A similar strategy was employed to create the *A. baumannii* AB5075 Aar seed region mutant strain (Aar*). This involved amplifying a ∼2,000 bp amplicon that was composed of the region upstream of *aar* and a portion of *aar* using primers containing the *aar* seed region mutation (primers 1 & 11). Similarly, a ∼2,000 bp amplicon composed of the region downstream of *aar* and a portion of *aar* was amplified using primers containing the *aar* seed region mutation (primers 4 & 12). A ∼4,000 bp chimeric PCR product encoding the point mutation (5′-CTCC-3′ → 5′-GTGG-3′) was then constructed by overlap extension PCR (Primers 1 & 4). The *A. baumannii* Δ*aar::sacB_aacC4* insertion mutant strain was then transformed with this PCR product to obtain Aar*, a strain with the mutated form of *aar*. The mutation was confirmed by Sanger sequencing (Eurofins Genomics).

### Plasmid constructions

PCR products were routinely purified using Monarch® PCR & DNA clean-up kit (NEB), or E.Z.N.A.® Cycle Pure Kit (Omega Bio-Tek) according to the manufacturers’ manuals. Plasmids were routinely purified using the E.Z.N.A.® Plasmid DNA Mini Kit I (Omega Bio-Tek). Plasmid constructions were carried out in *E. coli* TOP10 cells. The sequence of all inserts of constructed plasmids were confirmed by Sanger sequencing. To overexpress T4 RNA ligase, the *t4rnlI* gene was amplified by PCR using purified T4 phage lysate as a template (primers 13 & 14) and cloned into pVRL2Z using restriction digestion with HindIII and SalI followed by ligation with T4 DNA ligase (Lucidi *et al*., 2018). The SLiCE cloning procedure was used to construct the pP_L_, pXG10sf plasmids and pWH1266 derivative plasmids used in this study (Hunger *et al*., 1990; Urban and Vogel, 2007; Zhang *et al*., 2012; Corcoran *et al*., 2012). All DNA oligonucleotides used to amplify insert regions were modified at the 5′- and 3′-ends to include ∼20 base-pairs of homologous sequence to the ends of the linearised plasmid backbone created by PCR. The amplified linear plasmid backbone was DpnI-digested with Fastdigest DpnI (Thermo Fisher) to remove residual template DNA prior to the SLiCE reaction. To construct the pP_L_-Aar plasmid, the *aar* locus, from predicted TSS and including ∼150 nt downstream of the predicted terminator, was amplified (primers 15 & 16). This insert was cloned into the PCR-amplified pP_L_ plasmid backbone (primers 17 & 18) using SLiCE. The pP_L_-Aar* mutant was created by first amplifying ∼250 nucleotides upstream and downstream of the predicted Aar seed region primers designed to include the point mutations (19 & 11 and 10 & 12). An overlap extension PCR was then used to create a chimeric PCR product to act as a template for amplifying the pP_L_-Aar insert as before (primers 19 & 10). To construct the pXG10sf-target plasmids, inserts composed of the mRNA targets (including the TSS and predicted interaction site were amplified (primers 20 – 25). These inserts were then cloned into the linearised PCR amplified pXG10sf backbone (primers 26 & 27). A compensatory mutation in *carO* (*carO\**) was created in a similar manner to Aar* (primers 28 – 31). This was used as a template to construct pXG10-CarO*-sfGFP. To construct the pWH1266-mRNA-sfGFP plasmids, the mRNA target-sfGFP translational fusions from their respective pXG10sf plasmids were amplified (primers 32-34) and cloned into the linearised PCR-amplified pWH1266 backbone (primers 35 & 36). These constructs were placed under control of the β-lactamase (*bla*) promoter driving constitutive expression of the translational fusions.

### Hi-GRIL-seq experiment

*A. baumannii* AB5075 pVRL2Z-*t4rnlI* was streaked on L-agar containing zeocin and incubated at 37°C overnight. The next day, 5 mL L-broth containing zeocin was inoculated with a single, opaque (VIR-O) colony and grown over night (16 h, 37°C, 220 rpm, (Chin *et al*., 2018)). The next morning, the culture was diluted 1:1,000 into fresh 25 mL L-broth containing zeocin in a 250 mL Erlenmeyer flask and incubated at 37°C and 220 rpm agitation. After cells reached an OD600 of 2.0, 100 mM L-arabinose was added for one hour to induce expression of T4 RNA ligase before transcription was stopped by the addition of 2/5 of the culture volume of ice-cold “stop solution” (95% ethanol, 5% phenol) (Tedin and Bläsi, 1996). The imipenem and dipyridyl shocks were carried out by addition of imipenem (16 μg/mL, final concentration) or 2,2′-dipyridyl (0.2 mM, final concentration) for 10 min before T4 RNA ligase induction, stopping transcription as above and RNA isolation as described below. For the T4 RNA ligase “non-induced” control, cells were grown to OD 2.0 and incubated for an additional hour (without the addition of L-arabinose). The experiment was repeated independently to obtain a biological replicate.

### RNA extraction and RNA-sequencing

Total RNA for Hi-GRIL-seq and Northern blotting was isolated using TRIzol as described previously (Kröger *et al*., 2018) and was sent to Vertis Biotechnology AG (Freising, Germany) for DNase digestion, cDNA library preparation and sequencing as follows: Ribosomal RNA (rRNA) was depleted using a Vertis Biotechnology AG in-house developed depletion probes. The RNA was then fragmented using ultrasound (four pulses of 30 s at 4℃). An oligonucleotide adapter was ligated to the 3′ end of the RNA and first strand synthesis was performed with M-MLV reverse transcriptase and the 3′ adapter as the primer. The first strand cDNA was purified, and the 5′ Illumina TruSeq sequencing adapter was ligated to the 3′ end of the antisense cDNA. The resulting cDNA was amplified by PCR to 10-20 ng/μl using high fidelity DNA polymerase for 12-13 cycles. The cDNA was purified using Agencourt AMPure XP kit (Beckman Coulter Genomics) and analysed by capillary electrophoresis on a Shimadzu MultiNA microchip. For Illumina NextSeq sequencing, the samples were pooled in equimolar amounts and cDNA was size-fractioned (200-500 bp) using a preparative agarose gel. Finally, the cDNA pool was sequenced on an Illumina NextSeq 500 machine (75 bp read length).

### Bioinformatic analysis of Hi-GRIL-seq data

The quality of sequencing reads was assessed using FastQC. Short reads were first trimmed using trim_galore with default settings (https://www.bioinformatics.babraham.ac.uk/projects/trim_galore/) and then mapped against the *A. baumannii* reference genome including the three plasmids (NZ_CP008706.1, NZ_CP008707.1, NZ_CP008708.1, NZ_CP008709.1) containing manually added sRNA annotations using bowtie2 version 2.4.2 with the ’--very-fast’ option (Langmead and Salzberg, 2012; Kröger *et al*., 2018). Reads that could not be aligned were used for chimera detection. Sequences of 20 bp length were extracted from start and end of these reads and mapped separately against the reference genome using bowtie2 with the ’--very-sensitive’ option. Reads for which both start and end sequences could be aligned were annotated with overlapping genome features. Chimera RNA candidates were selected from reads where one of the ends overlapped with an sRNA and the other with a different annotated feature. The whole analysis process has been implemented as a series of Perl scripts, which can be accessed at https://github.com/khokamp/chimera. Mapping statistics can be found in **Supplementary Table 2**. Transcripts per million (TPM) values were computed using the formula presented by Li *et al*. ((Li *et al*., 2010), **Supplementary Table 3**). To reduce identification of transient sRNA-RNA interactions, chimeras from all growth conditions were merged and sRNA-containing chimeras with under ten sequencing reads were excluded from the analysis. Similarly, sRNA-containing chimeras with more than 10% of their sequencing reads derived from the non-induced control group were excluded from further analysis. Figures describing the Hi-GRIL-seq results were prepared using Integrated Genomics Viewer (IGV, (Robinson *et al*., 2011)) and circlize package on R (Gu *et al*., 2014).

### Sequence conservation analysis

10,998 draft *Acinetobacter* spp. genomes were downloaded from RefSeq (O’Leary *et al*., 2016). BLASTN version 2.14.1+ searches of the *A. baumannii* and *A. baylyi aar* sequences were performed against these draft genomes using default settings (Camacho *et al*., 2009). Lower percentage identity hits were filtered out to determine which regions of *aar* were subject to sequence divergence. A multiple sequence alignment of combined *A. baumannii* and *A. baylyi* hits was performed using Clustal Omega version 1.2.4 (Sievers *et al*., 2011). Jalview version 2.11.2.7 was then used to view this alignment and to cluster the samples using average distance (Waterhouse *et al*., 2009). Visualisation of these clusters was then performed using iTOL version 6.8.1 (Letunic and Bork, 2021). The Jalview conservation score was then exported as a text file and R was used to generate a conservation plot (R Core Team (2023) (Wickham *et al*., 2019)).

### Northern blotting

Northern blotting was performed using the DIG Northern Starter Kit (Roche) with digoxygenin (DIG)-labelled riboprobes as previously described (Kröger *et al*., 2012). In brief, DIG-labelled riboprobes were generated through *in vitro* transcription with T7 RNA polymerase using a PCR-generated template (primers 37 & 38). Five to 10 μg of total RNA were separated on a 7% (v/v) polyacrylamide gel (7.3 M urea, 1× TBE) and compared to RiboRuler LowRange RNA ladder (ThermoFisher Scientific). RNA from the gel was then blotted on a nylon membrane (Roche) at 125 mA at 4°C for 30 min. The RNA was crosslinked with UV light to the nylon membrane at 120 mJ. The membrane with the ladder was cut, stained in 2% methylene blue and de-stained in sterile dH_2_O at room temperature. The remainder of the membrane was processed according to the manufacturer’s description but with a 2 h blocking step. The ladder was reattached to the membrane before chemiluminescent imaging using an ImageQuant LAS4000 imager (GE Healthcare).

### *In vitro* transcription and radiolabelling of RNA

Radiolabelled RNA was prepared by *in vitro* transcription as previously described (Ryan *et al*., 2020). T7 promoter encoding primers were used to amplify templates from genomic DNA (primers 39 – 48), as outlined in **Supplementary Table 1**. The Aar* mutant was constructed by using the formerly mentioned overlap extension PCR product as a template, resulting in the substitutions (CUCC → GUGG, positions 29-32). *In vitro* transcription was performed using the MEGAscript T7 kit (Invitrogen) and template DNA removed with DNase I (1 U at 37°C for 15 min) prior to electrophoresis on a 6% (v/v) polyacrylamide gel (PAA) (7M urea, 1× TBE). *In vitro* transcription products were compared to a LowRange RNA ladder (ThermoFisher Scientific) and excised from the gel. The gel fragments were then suspended overnight in RNA elution buffer (0.1 M NaOAc, 0.1% SDS, 10 mM EDTA) at 8°C and shaking at 1,400 rpm. The eluted RNA was precipitated in ethanol:NaOAc (30:1), washed with 75% ethanol and eluted in 20 µL DEPC-treated water (Fisher). The *in vitro* transcribed RNA (50 pmol) was then dephosphorylated with 25 U of calf intestine alkaline phosphatase (Invitrogen) at 37°C for 1 h and extracted with phenol:cholorform:isoamylalcohol (P:C:I, 25:24:1). The dephosphorylated RNA (20 pmol) was then 5′-labelled (20 µCi of 32P-γATP) with 1 U of polynucleotide kinase (NEB) for 1 h at 37 °C. Radiolabelled RNA was then run on a G-50 column (GE Healthcare) and extracted from a polyacrylamide gel as above.

### Electrophoretic mobility shift assays (EMSA)

EMSAs were performed in 10 µL reactions as previously outlined (Ryan *et al*., 2020). Briefly, 5′-labelled Aar RNA (4 nM) was incubated for 1 h at 37°C with 1× structure buffer (Ambion), 1 µg yeast RNA and putative mRNA interaction partners at the indicated concentrations (0 nM, 16 nM, 32 nM, 62.5 nM, 125 nM, 250 nM, 500 nM and 1000 nM). Following incubation, 3 μL of 5× native loading dye (0.5× TBE, 0.2% (w/v) bromophenol blue, 50% (v/v) glycerol) was added to each reaction tube. These mixtures were then loaded on a 6% native PAA gel and electrophoresed in 0.5× TBE buffer at 4°C for 3 hours at 300V. Gels were then dried on Whatman paper and exposed to a phosphor screen, overnight. The phosphoscreen was imaged using a phosphorimager (TyphoonTM FLA 7000, GE Healthcare).

### In-line probing

In-line probing was performed by incubating 5′-labelled Aar (0.2 pmol) in 1× in-line probing buffer (50 mM Tris-HCl, pH 8.3, 100 mM KCl, 20 mM MgCl_2_) with the indicated concentrations of putative mRNA targets (0 pmol, 0.2 pmol and 2 pmol) at room temperature for 40 h as previously outlined (Sharma *et al*., 2007; Ryan *et al*., 2020). The reactions were then stopped with 10 µL 2× colourless gel-loading solution (10 M urea, 1.5 mM EDTA, pH 8.0). The Aar control ladder was prepared by mixing radiolabelled Aar (0.2 pmoL) with 10 µL 2× colourless gel-loading solution. The RNase T1 ladder was prepared by denaturing 0.4 pmol of 5′ labelled Aar with 8 µL of 1× sequencing buffer (Ambion) at 95°C for 1 min. This was then cooled at 37°C for 5 min before adding 1 µL of RNase T1 (0.1 U/µL) and further incubated at 37°C for 5 min. The alkaline hydrolysis ladder was prepared by incubating 0.4 pmol Aar with 9 µL alkaline hydrolysis buffer (Ambion) and incubated at 95°C for 5 min. Both ladders were mixed with colourless gel-loading solution prior to loading. All samples and ladders were then loaded and resolved on a 10% polyacrylamide gel (7M urea). Following this, the gel was dried, exposed to a phoshorscreen, and imaged using a phosphorimager as was performed for EMSAs.

Confirmation of RNA-RNA interactions in *E. coli*.

A plasmid constitutively expressing Aar (pP_L_-Aar) was constructed and the effect of Aar on target mRNA-sfGFP translational fusions (pXG10sf-*carO*, pXG10sf-*bfnH*, pXG10sf-*ABUW_RS07310*) was assessed by measuring GFP-mediated fluorescence compared to the fluorescence of strains co-expressing a control plasmid (pJV300) (Urban and Vogel, 2007; Corcoran *et al*., 2012). The translational fusions contained the predicted interaction sites, including the 5′ UTR and the first several codons of the mRNA targets *(carO* n=14, *bfnH* n =17 and *ABUW_RS07310* n=6 codons). Single colonies of *E. coli* carrying pJV300/pP_L_-Aar with pXG10-target plasmids were inoculated in 5 mL L-broth containing ampicillin and chloramphenicol. Dilutions (1:100) of overnight cultures of these strains were prepared and aliquoted in 96-well plates. Fluorescence (arbitrary units) and optical density at 600 nm (OD_600_) were measured in a Synergy H1^TM^ Hybrid Multi-Mode microplate fluorometer (BioTek) over 24 hours at 37°C with continuous orbital shaking and excitation (485 nm) and emission (508 nm). The fluorescence of different strains was compared by quantifying the fluorescence observed relative to optical density at 600 nm (fluorescence/OD_600_).

### Confirmation of RNA-RNA interactions in *A. baumannii*

Single colonies of wild-type, Δ*aar* and *aar*\* strains of *A. baumannii* AB5075 carrying the pWH1266 or pWH1266-mRNA-sfGFP plasmids were inoculated in 5 mL L-broth containing tetracycline. The next day, the culture was diluted 1:1,000 into fresh 25 mL L-broth containing tetracycline (12 μg/mL) in a 250 mL Erlenmeyer flask and incubated at 37°C and 220 rpm agitation for 16 h. The cultures were then aliquoted in 96-well plates. Fluorescence (arbitrary units) and optical density (OD_600_) were measured in a Synergy H1^TM^ Hybrid Multi-Mode microplate fluorometer (BioTek) with excitation (485 nm) and emission (508 nm). The fluorescence of different strains was compared by quantifying the fluorescence observed relative to optical density at 600 nm (fluorescence/OD_600_).

## RESULTS

### Establishment of Hi-GRIL-seq in *A. baumannii* AB5075

To uncover RNA interaction partners of sRNAs in *A. baumannii* AB5075 on genomic scale, we established Hi-GRIL-seq—a proximity ligation method that relies on transient overexpression of T4 RNA ligase to covalently attach interacting (i.e., base-pairing) RNA molecules in the cell (Zhang *et al*., 2017). The Hi-GRIL-seq method was originally developed in *P. aeruginosa* (Zhang *et al*., 2017) and its adaptation to *A. baumannii* AB5075 required several modifications to the protocol. Most importantly, the T4 RNA ligase was placed under control of a P_BAD_ promoter and ectopically expressed from a plasmid capable of replicating in *A. baumannii* AB5075 (pVRL2Z) (Lucidi *et al*., 2018). Dose-response experiments suggested 100 mM of L-arabinose as the optimal inducer concentration for efficient RNA-RNA ligation, while minimising lethality of the *A. baumannii* AB5075 culture (**Supplementary Figure 1A**). To perform Hi-GRIL-seq in *A. baumannii* AB5075, we grew cultures in L-broth to early stationary phase (OD_600_ 2.0) before inducing the ectopic expression of T4 RNA ligase for one hour **(Figure 1A**, IND). As a control, cells were grown to OD_600_ 2.0 in L-broth and incubated for an additional hour in the absence of L-arabinose (“non-induced”, NI). Because the expression of small regulatory RNAs is often activated in stressful environments (Holmqvist and Wagner, 2017), we additionally subjected these T4-induced cultures to brief (10 min) *in vivo*-relevant stress conditions. Specifically, in light of carbapenem-resistant *A. baumannii* posing a significant risk to global health (Tacconelli *et al*., 2018; Hamidian and Nigro, 2019), we exposed *A. baumannii* for 10 minutes to the beta-lactam antibiotic imipenem (IMIP). Alternatively, we limited iron availability by adding the iron chelator 2,2′-dipyridyl (DIP) to mimic the deprivation of infection-relevant iron (Cook-Libin *et al*., 2022). In any case, total RNA from biological duplicates was isolated and used to construct cDNA libraries for RNA-seq.

**Figure 1:**
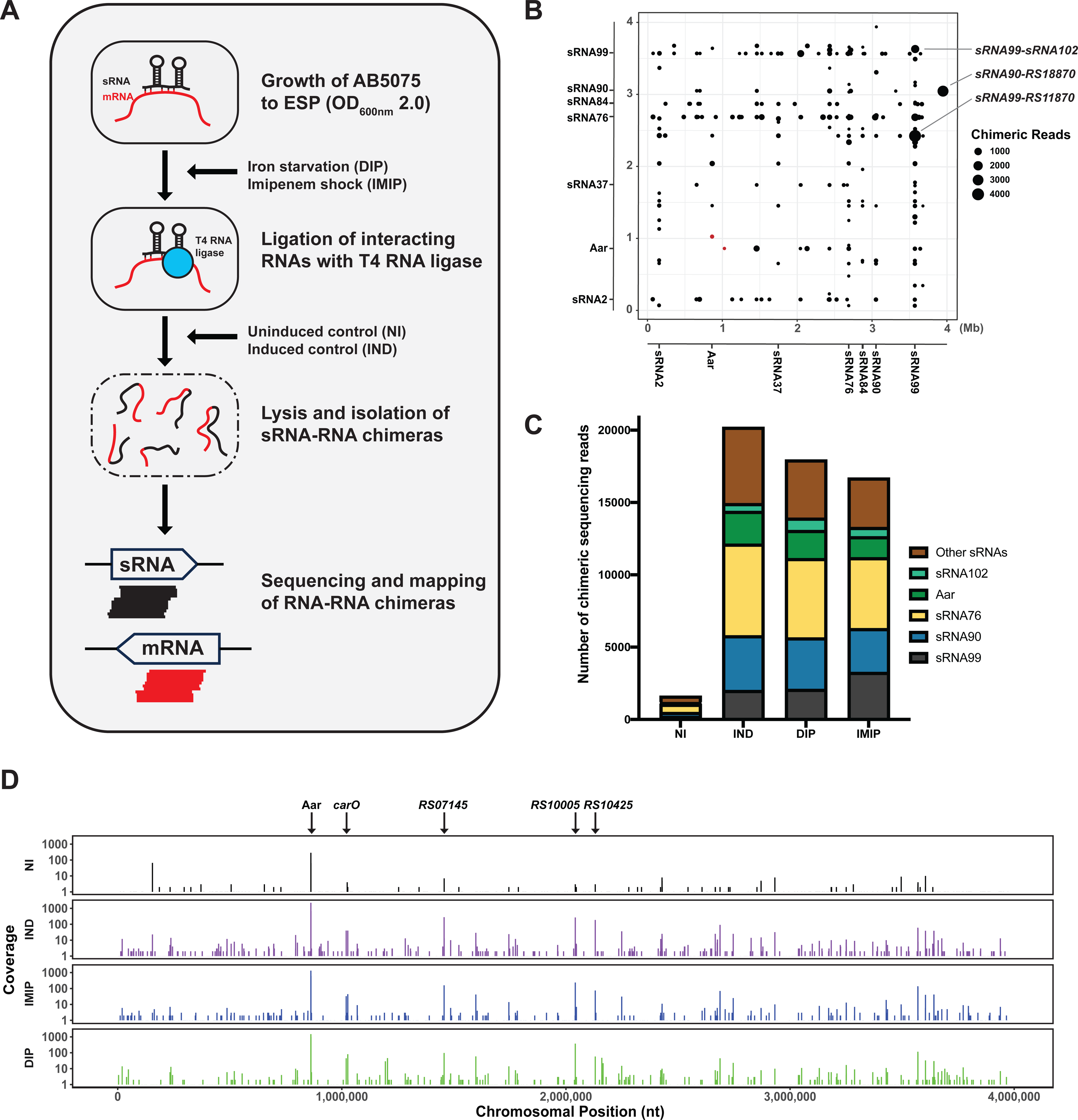
The proximity ligation experiment Hi-GRIL-seq was used to identify sRNA-mRNA interactions in *A. baumannii* AB5075. (**A**) Schematic depicting Hi-GRIL-seq workflow. **(B)** Global identification of sRNA-RNA interactions identified by Hi-GRIL-seq in *A. baumannii* AB5075 in the T4 RNA ligase induced (IND) control. The genomic locations of sRNA-containing chimeric reads were mapped to the AB5075 chromosome, where the x-axis represents the 5′ end of each chimeric fragment while the y-axis represents the 3′ end of the fragment. Only chimeric reads with ≥ 50 sequencing reads are shown. Each dot represents the exact genomic coordinate of these fragments. The size of the dot is proportional to the number of sequencing reads observed. The locations of the Aar-carO chimeric sequencing reads are labelled in red. **(C)** Proportion of sRNAs in RNA-RNA chimeras. The number of chimeric sequencing reads containing sRNA candidate sequences is shown for each condition, including non-induced (NI), induced (IND), iron starvation (DIP) and imipenem shock (IMIP). These chimeric reads are the combination of the biological duplicates of each treatment condition. **(D)** Mapped reads of Aar-containing chimeric sequencing reads across the *A. baumannii* AB5075 chromosome. Mapped reads were prepared by combining the chimeric sequencing reads from treatment condition replicates. Lines indicate the location and abundance of the chimeric junctions. The most abundant mRNA interaction partners ligated within sRNA-containing chimeras are indicated.

### Analysis of RNA-RNA chimeras to identify candidate regulatory sRNA-mRNA interactions

Based on comparable datasets from *P. aeruginosa* (Zhang *et al*., 2017), we expected only a small fraction of the sequence reads to be chimeric. We therefore opted for a sequencing depth beyond a typical bacterial transcriptomic experiment and generated a total of >77 million sequence reads per library (**Supplementary Table 2**). We mapped the Hi-GRIL-seq reads to the *A. baumannii* AB5075 genome and calculated the abundance of reads mapping to, and outside of, annotated features (**Supplementary Table 2, Supplementary Figure 1B**) and calculated transcripts per million (TPM) values (**Supplementary Table 3**). In all conditions and replicates, more than 82% of the reads mapped uniquely to the genome, indicating that these reads were non-chimeric (**Supplementary Table 2**). To identify chimeric reads, the first and last 20 nucleotides of each unmapped read was separately aligned to the AB5075 genome. On average, there was a higher percentage of chimeric reads (0.9-1.4%) in the IND, IMIP and DIP conditions compared to the NI control (0.8-0.9%, **Supplementary Table 2**), however, obtaining chimeric RNA reads in the non-induced samples indicates that the pBAD promoter expressed basal levels of T4 RNA ligase. Of the chimeric reads, 3.5-5.5% from the IND, DIP and IMIP conditions contained an annotated sRNA on one end and another genomic feature at the other end and were consequently considered as candidate sRNA-RNA interaction partners (**Figure 1B, Supplementary Figure 2**). In those sRNA-RNA chimeras, we detected 86 sRNAs of AB5075.

*In vivo* RNA proximity ligation can result in the ligation of RNA molecules that are only transiently in close physical vicinity and do not entail any functional consequences. To increase the chances of identifying functional base-pairing events, sRNA-containing chimeras that were scarce (under 10 sequencing reads) or were overrepresented in the non-induced condition (more than 10% of reads) were excluded from further analysis. Applying these cut-offs reduced the number from 86 to 40 sRNAs contained in sRNA-RNA chimeras. We identified 632 distinct chromosomal sRNA-containing chimeras and 74 sRNA-containing chimeras that were ligated to plasmid-derived RNAs (**Supplementary Figure 3 and Supplementary Table 4**). The sRNAs sRNA76, Aar (sRNA21), sRNA90 and sRNA99 accounted for 30.6%, 10.2%, 18.9% and 14.4% of chimeric reads, respectively, suggesting that they represent “keystone sRNAs” that form regulatory hubs in *A. baumannii* (**Figure 1C**, (Gebhardt *et al*., 2023)). The sRNA-containing chimeric reads were distributed across the AB5075 chromosome, an example of this is shown for Aar (**Figure 1D**).

### Hi-GRIL-seq reveals Aar interaction partners including *carO* mRNA

To validate our high-throughput approach, we selected the conserved sRNA Aar for further investigation (**Figure 3A and Supplementary Figure 7A**). Aar was previously suggested to act as a posttranscriptional regulator of amino acid metabolism in *A. baylyi*, however, no direct Aar targets have been identified in *Acinetobacter* spp. (Schilling *et al*., 2010). Here, we found 98 different Aar- containing chimeras in the Hi-GRIL-seq data that met our cut-off parameters, suggesting that Aar may be a regulatory RNA that acts primarily through base-pairing (**Figure 2A, Supplementary Table S4**). Integrating the Hi-GRIL-seq data with IntaRNA *in silico* predictions based on hybridisation energy and location of the putative interaction sites (Busch *et al*., 2008), suggested that Aar base-pairs with the ribosome binding sites (RBS) of *carO*, *bfnH* and *ABUW_RS07310* mRNAs and within the “5-codon window” of *ABUW_RS04325* mRNA (**Figure 2B & C** and **Supplementary Figure 4**). Four Aar nucleotides (positions 29-33, 5′-CUCC-3′) were present in all predicted interactions, suggesting that they might be vital for sRNA binding and act as the Aar “seed” region that initiates contact with the target molecules (**Figure 2B & C**).

**Figure 2:**
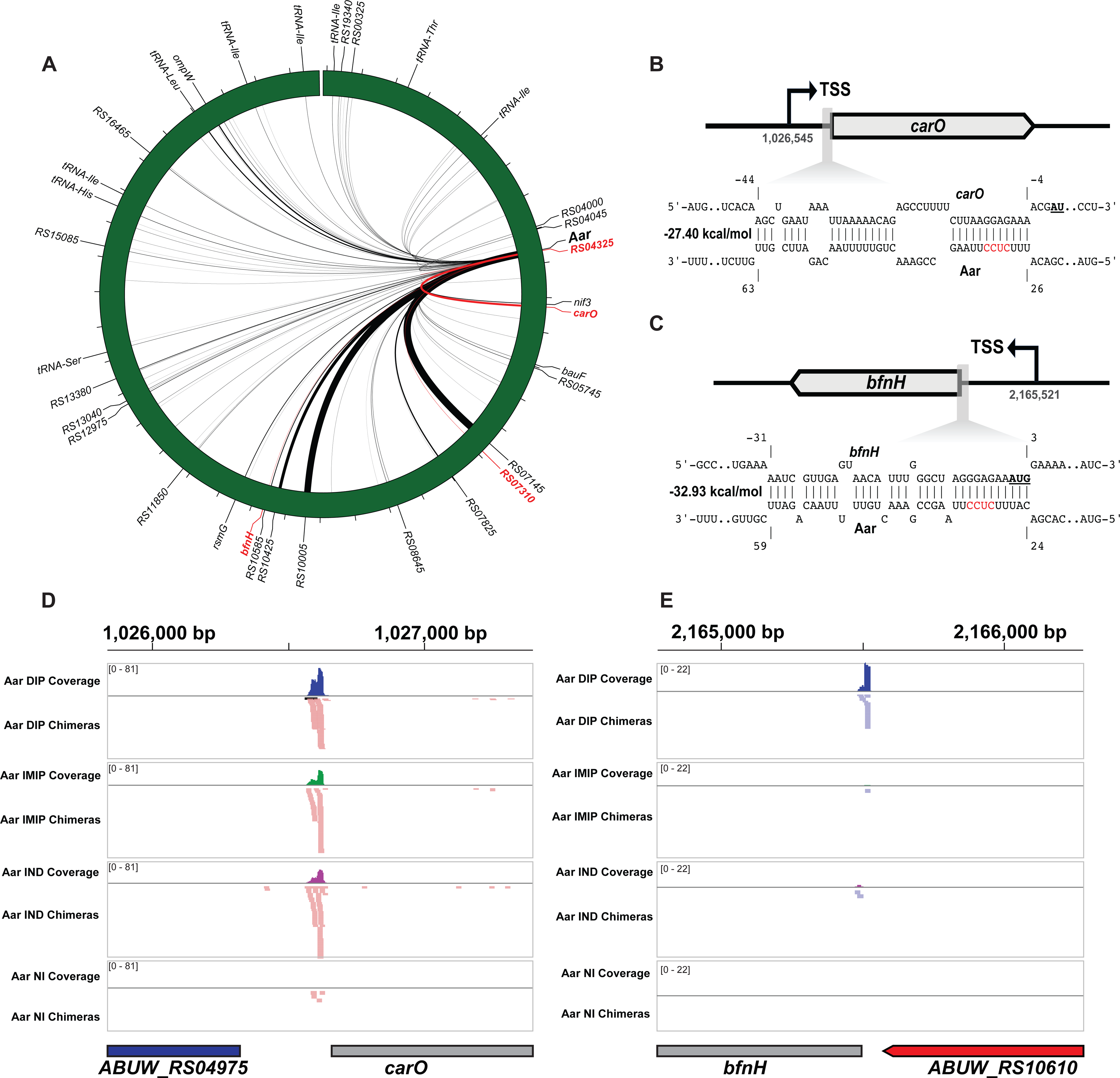
Hi-GRIL-seq reveals target molecules of sRNA Aar. (**A**) Hi-GRIL-seq revealed that the *A. baumannii* sRNA Aar was found to interact with other RNA molecules (n=88). These interactions are shown across the chromosome where the thickness of the line is proportional to the number of chimeric reads obtained. Aar interaction partners studied in detail are outlined in red. **(B & C)** The sRNA-mRNA interaction prediction tool IntaRNA was used to identify likely interactions between Aar and putative mRNA targets (Busch *et al*., 2008). The position of the IntaRNA predicted interaction duplexes with *carO* (**B**) and *bfnH* (**C**) mRNAs are shown. These interactions are predicted to sequester the ribosome binding site (RBS). The chromosomal positions of the transcriptional start sites (TSSs) are shown below the curved arrow. The start codon of both mRNA targets is highlighted in bold and underlined. The location of the predicted interaction is relative to these start codons. The predicted Aar seed region is highlighted in red. **(D & E)** The location of the *carO* (**D**) and *bfnH* (**E**) portions of chimeric fragments (*aar*-*carO* and *aar*-*bfnH,* respectively) were mapped to the AB5075 chromosome. The location of these chimeric reads is visualised for each condition used in Hi-GRIL-seq. A coverage profile of the number of chimeric reads is shown on the upper part of each track. Individual chimeric reads are shown in the lower part of each track.

**Figure 3:**
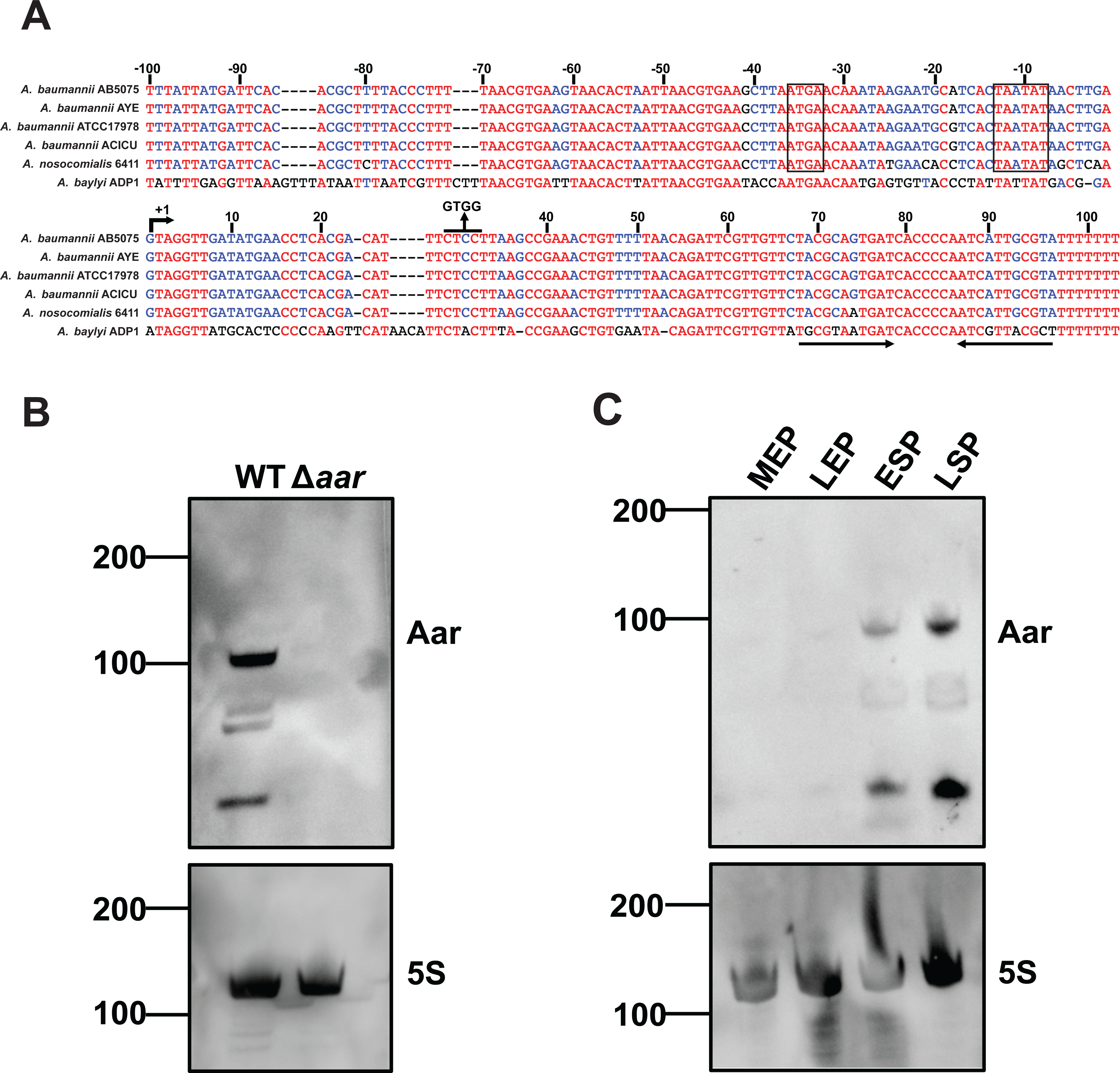
Aar is a growth-phase dependent sRNA that is conserved among pathogenic *Acinetobacter* species. (**A**) Sequence alignment of *aar* across several *Acinetobacter* species. This sequence alignment was performed using the Multalin tool, the colour of nucleotides corresponds with the extent of conservation. The predicted transcriptional start site (TSS) is indicated by +1. The numbers above the sequence represent the distance from the TSS. A line is drawn above the predicted seed region and its mutated form is indicated. The putative –10 and –35 regions are indicated with a box (for *A. baumannii* and *A. nosocomialis* 6411). The Rho-independent terminator of Aar is shown by two arrows below the sequence. (**B**) Northern blotting of Aar in *A. baumannii* AB5075 wild-type and Δ*aar* strains. RNA was isolated from LSP. Numbers depict sizes of RNA ladder in nucleotides. (**C**) The expression of Aar across growth of *A. baumannii* AB5075 as determined by Northern blotting. RNA was isolated from *A. baumannii* grown to MEP, LEP, ESP and LSP. Numbers depict sizes of RNA ladder in nucleotides. Expression of Aar and the loading control 5S rRNA were detected using their respective DIG-labelled riboprobes. Three independent biological replicates were performed for each Northern blotting experiment and one representative blot shown.

Examining the exact genomic locations of individual chimeric reads for the four putative mRNA targets (**Figure 2D & E and Supplementary Figure 6**), we found that 242 (∼86%) of the identified *carO* chimeric reads mapped to the predicted interaction site (combination of replicates from all conditions), relatively equally distributed across all experimental conditions (**Supplementary Figure 5A)**. Twenty-four (∼86%) of *bfnH* chimeric reads mapped to the predicted interaction site (from all conditions), mostly contributed by the iron starvation condition (**Figure 2E, Supplementary Figure 5B**) in line with this gene’s induction in response to iron chelation (Eijkelkamp *et al*., 2011; Nwugo *et al*., 2011). Similarly, 19 (∼53%, from all conditions) and twelve (∼92%, from all conditions) of *ABUW_RS07310* and *ABUW_RS0432*5 chimeric reads mapped to their IntaRNA predictions, respectively, relatively homogeneously dispersed across the four different experimental conditions (**Supplementary Figure 5C & D**).

Together, the mapping of chimeric reads to the RBS and the 5-codon window suggests that Aar may act as a translational regulator of these mRNA molecules. A web-based browser is available at: http://bioinf.gen.tcd.ie/jbrowse2/Hi-GRIL-seq showing the mapped reads from Hi-GRIL-seq in addition to coverage plots of the Aar-containing chimera across the chromosome and plasmids.

Aar is a ∼100 nt small RNA conserved in pathogenic *Acinetobacter* spp. that accumulates in stationary phase in *A. baumannii* AB5075.

While Aar is solely present among *Acinetobacter* species, the sRNA displays only limited sequence conservation between *A. baylyi* ADP1 compared to four model *A. baumannii* strains and *A. nosocomialis* 6411 (**Figure 3A**). The different sizes of Aar between *A. baumannii* (103 nt, (Kröger *et al*., 2018)) and *A. baylyi* (181 nt, (Schilling *et al*., 2010)) may be attributed to the use of different transcriptional start sites (TSSs). The *A. baumannii aar* promoter architecture bears striking resemblance to the consensus promoter in *A. baumannii* ATCC17978, suggesting that Aar expression may be dependent on the sigma factor RpoD (Kröger *et al*., 2018). To perform a more comprehensive analysis of *aar* conservation, we used a BLAST-based approach to 10,998 draft *Acinetobacter* spp. genomes. Of the 8,801 *aar* sequences identified in *A. baumannii*, the majority (7,940) had 100% sequence identity with the corresponding AB5075 homologue. We retained only the lower percentage identity matches and then performed a multiple sequence alignment of the sequences matching *A. baumannii* and *A. baylyi aar* to understand which regions of the gene were subject to sequence divergence (**Supplementary Figure 7A**). Overall, *aar* appeared to have lower conservation at the 5′ end and greater conservation towards the 3′ end. Visualisation of the *aar* clusters revealed that the *A. baylyi* and *A. baumannii* homologues formed distinct groups (**Supplementary Figure 7B**). The *A. baylyi* group involved a subgroup containing shorter *aar* sequences. *In silico* predictions of the *A. baumannii* AB5075 Aar secondary structure suggests it to adopt a mostly single-stranded structure with a hairpin at the 5′ end and two hairpins at the 3′ end followed by a poly(U) tail to promote Rho- independent termination (**Supplementary Figure 8A**). The two hairpins at the 3′ end of Aar harbour the highest sequence conservation and might be central for functionality of the molecule.

To assess the size of Aar in *A. baumannii* AB5075, we employed Northern blotting using a riboprobe that spans the entire predicted length of the sRNA (**Figure 3B**). Bands of smaller size are apparent, which could represent partially degraded or processed Aar variants, albeit analysis of *aar* sequencing reads did not reveal any discernible processing sites (**Supplementary Figure 8B**). All bands disappeared upon deletion of *aar*, indicating that they emanate from the *aar* locus and that full-length Aar is a ∼100 nt transcript in *A. baumannii* as predicted earlier from dRNA-seq data (**Figure 3B**, (Kröger *et al*., 2018)). To investigate whether Aar displays differential expression during growth, we performed Northern blotting using total RNA isolated from four stages of growth in LB (mid-exponential phase (MEP, OD_600_ = 0.3, late exponential phase (LEP, OD_600_ = 1.0) early stationary phase (ESP, OD600 = 2.0) and late stationary phase (LSP, 16h of growth) (**Figure 3C**). We observed that Aar is expressed in a growth phase-dependent manner with higher expression in early and late stationary phase and low abundance in earlier growth stages.

To determine whether expression of Aar was altered in different environmental conditions, we exposed *A. baumannii* to multiple stressors before isolating RNA and performing Northern blotting. This involved growing *A. baumannii* AB5075 to LEP, when Aar expression is low (**Figure 3C**), at either 37°C or 25°C and shocking the culture for 15 min. The samples grown at 37°C were shocked with either 0.3 M NaCl, 200 µM 2,2′-DIP or 1 µg/mL polymyxin B to expose *A. baumannii* to osmotic shock, iron starvation or envelope stress, respectively. Samples that were grown at 25°C were transferred to 37°C for 15 min to act as a temperature shift (TS) from ambient to body temperature. Mock control groups were included, where the samples were not shocked but grown at their respective original temperature for 15 min. Aar was more abundant following osmotic shock (4-fold increase) and upon iron limitation (2-fold increase) (**Supplementary Figure 9**), which matched previous work in *A. baylyi* (Schilling *et al*., 2010). Surprisingly, unlike in *A. baylyi*, we found that *A. baumannii* Aar was more abundant when grown in ambient (25°C) than physiological (37°C) temperatures (2-fold increase) and appears to be induced following TS (2.6-fold increase). Addition of polymyxin B had no effect on Aar abundance, suggesting that Aar expression is not induced by envelope stress. Therefore, Aar expression is responsive to environmental changes like osmotic stress and temperature shifts, potentially allowing *A. baumannii* to fine-tune the expression of target molecules in these environments.

### Aar binds to *carO* and *bfnH* mRNAs *in vitro* using a conserved seed region

To investigate whether Aar interacts with *carO* and *bfnH* mRNAs *in vitro*, we performed electrophoretic motility shift assays (EMSAs). EMSAs were performed using radiolabelled Aar with increasing concentrations of ∼150 nt-long RNA molecules (from the transcriptional start site of the target molecules and including the predicted Aar binding site (**Figure 4 and Supplementary Figure 10**). These EMSAs revealed that Aar binds to increasing concentrations of *carO* and *bfnH*, resulting in band shifts with high affinity interactions (*K*_D_ = 55.78 nM and *K*_D_ = 85.86 nM, respectively) (**Figure 4A-D**). Aar was also found to form band shifts with increasing concentration of *ABUW_RS07310* (*K*_D_ = 42.8 nM) and *ABUW_RS04325* (*K*_D_ = 347.7 nM) (**Supplementary Figure 10A and B and Supplementary Figure 11A and B**). The interaction with *ABUW_RS04325* was notably weaker than with the other mRNA targets, suggesting that it may be less relevant in *A. baumannii* or that it depends on the presence of an RNA chaperone such as Hfq. To determine whether the predicted “seed” region mediated these base-pairing interactions, we mutated Aar from 5′-CUCC-3′ to 5′-GUGG-3′ (Aar*) and repeated the EMSAs with each target (**Supplementary Figure 10C-F**). Binding to the four target molecules was abolished when using Aar* (**Figure 4A and 4B and Supplementary Figure 11A** and **4B**) demonstrating that the 5′-CUCC-3′ four-nucleotide sequence is essential for RNA-RNA interactions.

**Figure 4:**
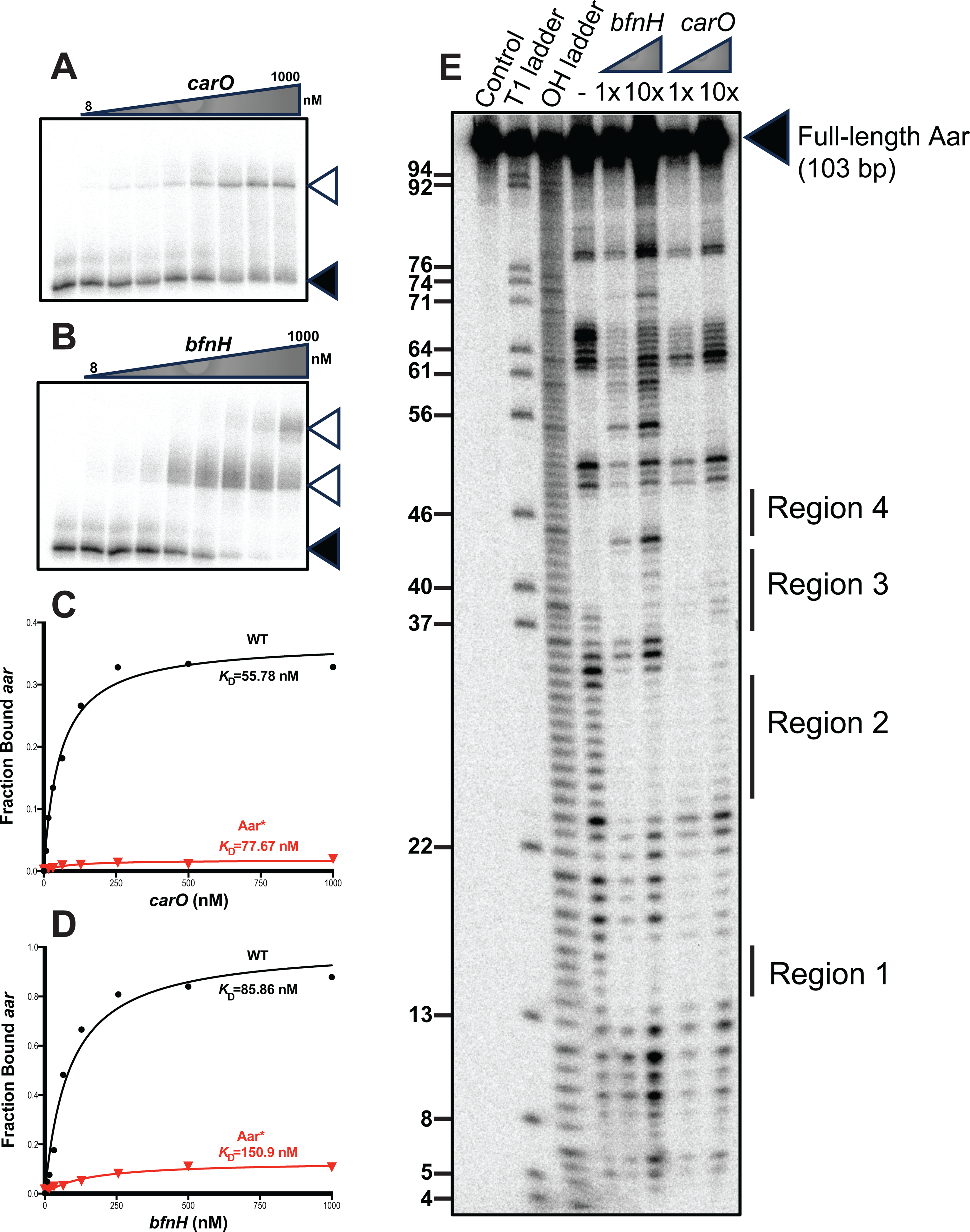
Aar interacts with *carO* and *bfnH* mRNAs using a common seed region. The ability of 5′-radiolabelled Aar to interact with increasing concentrations (from 8 nM to 1000 nM) of (**A)** *carO* and (**B)** *bfnH* was assessed using electrophoretic motility shift assays (EMSAs). This revealed that full length Aar (◄) formed discernible band shifts (◁) with increasing concentrations of these mRNA targets. EMSAs with radiolabelled wild-type Aar (WT, black) and seed region mutated Aar (Aar*) interacting with (**C**) *carO* and (**D**) *bfnH* were used to quantify the dissociation constants (*K*_D_) of these targets respectively. The *K*_D_ values were calculated from the mean of two independent replicates (n=2). (**E**) The exact Aar nucleotides involved in *bfnH* and *carO* base-pairing were identified using in-line probing. The protected regions (Region 1, Region 2, Region 3 and Region 4) are indicated. This was performed using three independent replicates (n=3) and one representative blot shown.

Next, we determined the secondary structure of Aar and the exact Aar nucleotides that base-pair with *bfnH* and *carO* mRNAs by employing in-line probing (**Figure 4E, Supplementary Figure 11**). When incubated *in vitro* in isolation, Aar adopts a single stranded 5′-end structure (up to nucleotide 38) followed by two hairpin loops (**Supplementary Figure 12A**). This experimentally inferred secondary structure of Aar largely matches the *in silico* prediction (**Supplementary Figure 8),** except for the predicted hairpin at the 5′-end, which is absent *in vitro*. In-line probing further revealed that two and four adjacent regions of Aar (termed Region 1 to 4) were involved in *carO* and *bfnH* base-pairing, respectively. Region 1 was protected in the presence of both the *carO* and *bfnH* 5′ region (positions 15-17, **Figure 4E, Supplementary Figure 12B**). Similarly, region 2, encompassing the predicted seed region, was protected from cleavage in the wildtype Aar when incubated in the presence of *bfnH* (positions 25-33) and *carO* (positions 25-38) mRNA **(Figure 4E, Supplementary Figure 12)**. Disruption of the predicted seed region (visible at positions 29, 31 and 32 of the T4 ladder) in the Aar* mutant variant resulted in a loss of cleavage protection at region 2 (**Supplementary Figure 11C**). Furthermore, the incorporation of these mutations did not affect the local secondary structure around the seed region *in vitro*, demonstrating the importance of this sequence in initiating inter-molecular base-pairing. In addition, binding to *bfnH* mRNA resulted in a structural rearrangement of Aar, where the first hairpin loop of the sRNA unfolded into a single-stranded region (positions 54-61, **Figure 4E**) and additional base-pairing protected regions 3 and 4 (positions 37-43 and 45-48, respectively), resulting in an extensive Aar*-bfnH* duplex structure (**Supplementary Figure 12**). Together, these results illustrate that *in vitro*, Aar initiates base-pairing with its target mRNA molecules using a contiguous seed encoded within a single-stranded region near the 5′-end.

### Aar represses translation of *carO, bfnH* and *ABUW_RS07310* in a heterologous host

To probe Aar-mediated regulation *in vivo*, we employed an established two-plasmid fluorescent reporter system in *E. coli* (Urban and Vogel, 2007; Corcoran *et al*., 2012). This heterologous system was initially selected due to the lack of compatible plasmids that can co-replicate in *A. baumannii*. A similar strategy was employed in *Vibrio cholerae* to investigate sRNA-mRNA base-pairing (Huber *et al*., 2022). By measuring the relative fluorescence of superfolder-GFP (sfGFP) translational fusions with Aar target mRNAs (*carO*, *bfnH* and *ABUW_RS07310*) in the presence and absence of Aar, we determined the regulatory potential of this sRNA. Expression of Aar significantly reduced the fluorescence intensity of the BfnH-sfGFP (4.7-fold; p<0.0001), CarO-sfGFP (6-fold; p<0.0001) and ABUW_RS07310-sfGFP (1.5-fold; p<0.05) compared to the control strain, suggesting that Aar is inhibiting translation of these three targets (**Figure 5A and 5B, Supplementary Figure 13**). In the Aar* mutant strain, the repression was abolished, restoring fluorescence of CarO-GFP and partially restoring BfnH-sfGFP fluorescence levels to levels of the control strain. The partial restoration may be due to more extensive base-pairing between Aar and *bfnH* mRNA outside of region 2 (**Supplementary Figure 12**). We were unable to construct functional compensatory mutations of the target fusions, as substitution of the nucleotides complementary to the Aar seed region, would disrupt their Shine-Dalgarno regions, preventing translation altogether (**Figure 2C and Supplementary Figure 4 & 13**). In summary, these data suggest that Aar prevents translation by sequestering the Shine-Dalgarno sequence of at least three of its target mRNAs in a heterologous system.

**Figure 5:**
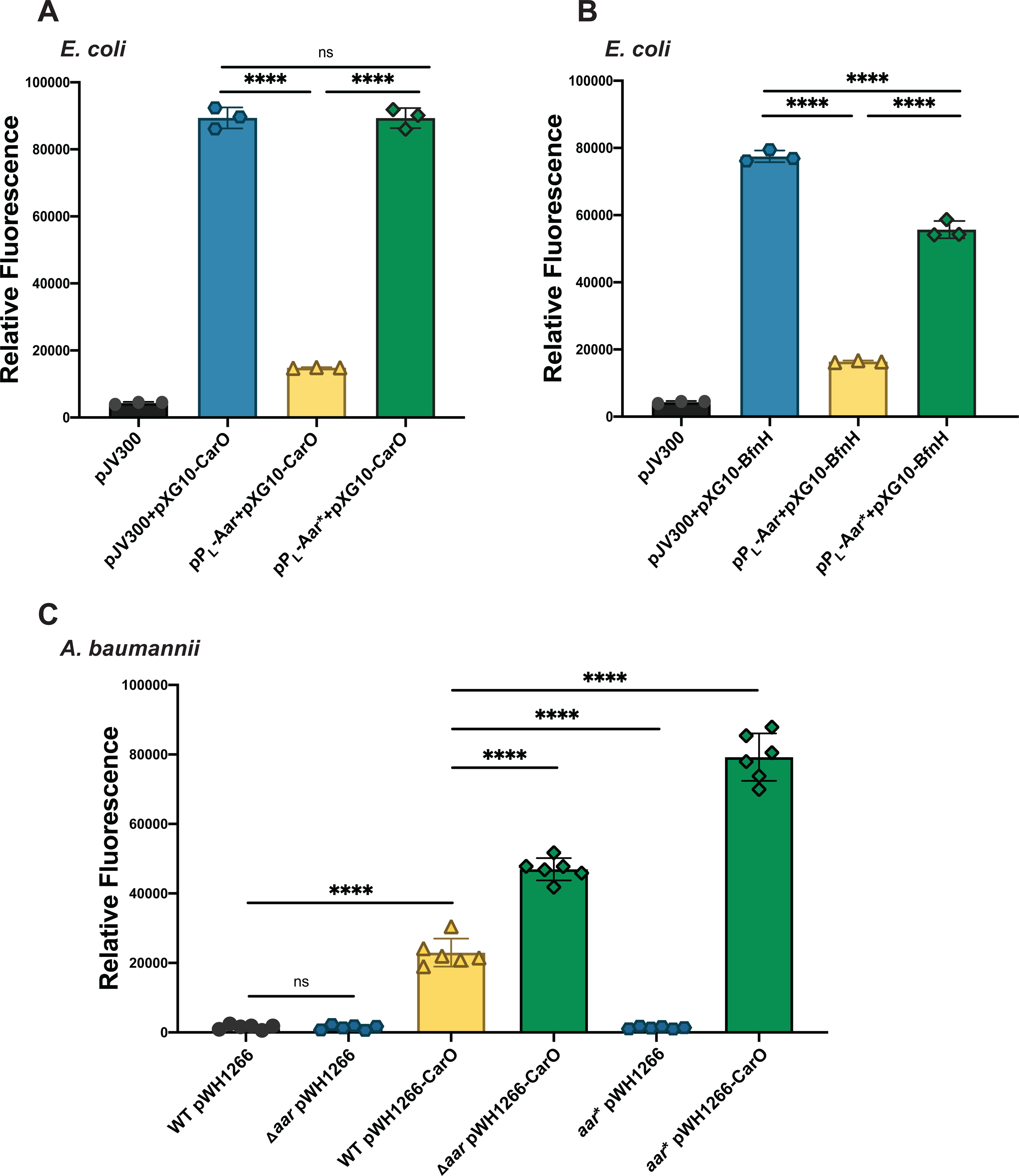
Translational reporters in *E. coli* and *A. baumannii* AB5075 reveal that Aar suppresses translation of *carO* and *bfnH* mRNAs using the conserved seed region. The involvement of Aar in modulating expression of CarO **(A)** and BfnH **(B)** was assessed using a two-plasmid reporter system in *E. coli*. The relative fluorescent intensity of strains carrying the pXG10-CarO/BfnH plasmid with either the control plasmid (pJV300), an Aar overexpression vector (pP_L_-Aar) or an Aar* overexpression vector (pP_L_-Aar*) was measured. A strain only carrying pJV300 was also included to measure autofluorescence of *E. coli*. Error bars represent the standard deviation from three independent biological replicates. Statistical comparisons were performed using one-way ANOVA followed by Tukey’s multiple comparisons test. **(C)** The involvement of Aar and Aar* in modulating expression of CarO was assessed in *A. baumannii* AB5075 using a translational reporter. The relative fluorescent intensity of wild-type strain (WT) and Aar-deletion (Δ*aar*) or Aar seed region mutant (*aar*\*) strains of *A. baumannii* AB5075 carrying the pWH1266-CarO plasmid was compared at LSP. Control strains carrying the pWH1266 plasmid were also measured to quantify the autofluorescence of strains. Error bars represent the standard deviation from six independent biological replicates. Statistical comparisons were performed using One-way ANOVA followed by Tukey’s multiple comparisons test. Differences were considered statistically significant where ns denotes P>0.05, * denotes P<0.05, ** denotes P<0.01, *** denotes P<0.001 and **** denotes P<0.0001.

### Aar represses CarO expression in *A. baumannii* AB5075 *in vivo*

To ensure that Aar-mediated repression of CarO and BfnH occurs in *A. baumannii,* we created CarO-sfGFP and BfnH-sfGFP translational reporters on the *A. baumannii*-*E. coli* pWH1266 shuttle plasmid under control of the β-lactamase promoter driving constitutive expression (Hunger *et al*., 1990). Fluorescence was measured at LSP in the wildtype and Δ*aar* genetic backgrounds. There was significantly higher relative fluorescence of CarO-sfGFP (2-fold; p<0.0001) in Δ*aar* than in wild-type AB5075, reaffirming the repressive role of Aar (**Figure 5C**). To demonstrate that the Aar seed region is vital for repression of CarO, we created an Aar seed region mutant strain of *A. baumannii* AB5075 (*aar*\*; 5′-CUCC-3′ to 5′-GUGG-3′ exchange of the seed region on the chromosome) and detected a significantly higher fluorescent intensity (2.9-fold; p<0.0001) in the *aar*\* strain than in the wild-type background (**Figure 5C**). This illustrates that physiological levels of Aar are sufficient to regulate CarO translation and that the sRNA’s seed region is critical for this activity. We did not detect a significant change in BfnH-sfGFP (p=0.2096) in neither the Δ*aar* nor the *aar*\* strain compared with the wildtype (**Supplementary Figure 14A, B**). The lack of BfnH repression in this translational reporter system may be attributed to the low endogenous levels of Aar and its lower affinity to *bfnH* than to *carO* mRNA (see **Figure 4C, D**). However, based on these data we conclude that at least CarO is not only repressed by Aar upon overexpression in a heterologous system, but also in its native *A. baumannii* environment and under physiological conditions, as was inferred from our global RNA-RNA interactome atlas.

## DISCUSSION

Over the last two decades, the field of sRNA-mediated gene regulation has exploded, driven by the precise identification of small, often non-coding transcripts through the application of deep-sequencing technologies (Sharma and Vogel, 2009; Colgan *et al*., 2017; Hör *et al*., 2018; Papenfort and Melamed, 2023). Following the identification of new candidate sRNAs (Holmqvist and Vogel, 2018), their functional characterisation and prediction of interaction partners is now also facilitated by deep-sequencing-based approaches. For example, several methods have recently been established that rely on RNA proximity ligation and subsequent sequencing of the generated RNA chimera (Hör *et al*., 2018; Melamed, 2020), albeit none of them has previously been applied in *A. baumannii*.

In this study, the RNA proximity ligation technique Hi-GRIL-seq that was originally developed in *P. aeruginosa* was optimized and applied to *A. baumannii* AB5075 (**Figure 1A** (Zhang *et al*., 2017)). The RBP-independent Hi-GRIL-seq method was deliberately chosen to expand our knowledge of the RNA biology of *A. baumannii*—a bacterium in which a global role of Hfq remains questionable. The strengths but also limitations of the Hi-GRIL-seq method became apparent during our investigation. For example, we observed that certain sRNAs, particularly sRNA2, were ligated in chimeras in the non-induced control condition but were not observed in induced samples (**Supplementary Figure 2**). This could be explained by the much higher abundance of sRNA2 (on average ∼5.5-fold higher compared to the other conditions) in the non-induced condition and by basal transcription of T4 RNA ligase from the pBAD promoter (**Supplementary Table 3**). This may have led to ligation of highly abundant sRNAs and non-base pairing RNA molecules. Alternatively, the high sequencing depth employed in this study was notably higher than those used for conventional bacterial RNA-sequencing, perhaps causing overrepresentation of transient, non-functional RNA-RNA interactions. Additionally, proximity ligation-based discovery tools exclusively capture interactions between monophosphorylated sRNAs and their RNA interaction partners, potentially excluding interactions mediated by triphoshorylated sRNAs (Han *et al*., 2016). We also noticed a lower overall percentage of chimeric reads containing sRNAs (∼0.06%) compared to the Hi-GRIL-seq study performed by Zhang and colleagues (0.26–0.29%, (Zhang *et al*., 2017)). This discrepancy might be due to differences in the experimental approach; e.g. Zhang and colleagues used an IPTG-inducible system to drive T4 RNA ligase expression, while we were reliant on induction using the pBAD promoter (Zhang *et al*., 2017) or by potentially different phosphorylation states at the 5′-ends of transcripts between *P. aeruginosa* and *A. baumannii*.

Despite these challenges, Hi-GRIL-seq allowed us to identify genuine sRNA-mRNA interactions in *A. baumannii* AB5075, yielding a total of 706 putative sRNA-RNA interactions for future targeted investigation. Other proximity ligation experiments have employed identification of sRNA seed regions and their corresponding binding sites to functionally characterise sRNAs (Melamed *et al*., 2016; Waters *et al*., 2017). Similarly, we focused here on the sRNA Aar, which is conserved among *Acinetobacter* species (**Figure 3A and Supplementary Figure 7A)**. Aar was initially speculated to be involved in *A. baylyi* amino acid metabolism, as transcripts of amino acid metabolic genes were differentially regulated in an Aar overexpression strain (Schilling *et al*., 2010). Here, we showed that—similar to its expression in *A. baylyi—*Aar expression was induced in *A. baumannii* following osmotic shock and iron limitation (**Supplementary Figure 9**). This is in spite of major differences in the *aar* promoter regions between the two species, which warrants further investigation as to whether Aar is regulated by common transcription factors in both organisms. While previous work in *A. baylyi* suggested that *aar* encodes a ∼180 nt-long molecule, we demonstrated that in *A. baumannii*, *aar* encodes a 103-nt sRNA, in agreement with differential RNA-seq data from this species (Schilling *et al*., 2010; Kröger *et al*., 2018). Furthermore, we predicted that a subcluster of *A. baylyi* strains encode a smaller variant of Aar, whose primary sequence is almost perfectly conserved (**Supplementary Figure 7A** and **B**). The greater conservation of this shortened *A. baumannii aar* allele among *Acinetobacter* species suggests that the functionally important sequence is located within this region, yet future studies may address whether different Aar isoforms have divergent roles between *A. baumannii* and *A. baylyi*.

Integration of Hi-GRIL-seq chimeric read information with *in silico* predictions of inter-molecular sequence complementarity suggested that Aar base-pairs with various mRNAs, most convincingly with the translation initiation regions of *carO*, *bfnH*, *ABUW_RS07310* and *ABUW_RS04325*, but none of them is predicted to be directly involved in amino acid metabolism as suggested for *A. baylyi* (Schilling *et al*., 2010). We identified a contiguous sequence of nine nucleotides within the Aar 5’ region that was present in all the predicted duplex structures with these four target mRNAs. This Aar seed region falls within a single-stranded region of the sRNA, rendering it readily available for inter-molecular base-pairing. Drawing from a series of *in vitro* biochemistry approaches, we confirmed these target binding models for *carO*, *bfnH* and *ABUW_RS07310*. To assess whether Aar binding entails any functional consequences for the expression of its target genes, we employed an established translational reporter system (Urban and Vogel, 2007) in *E. coli* and developed a new system in *A. baumannii*. In the heterologous system, overexpression of Aar repressed *carO*, *bfnH* and *ABUW_RS07310* translational fusions (**Figure 5A, 5B** and **Supplementary Figure 13A**). Furthermore, the disruption of the Aar seed region relieved repression of *carO* and *bfnH*. The use of a translational reporter system in wild-type and Aar-mutant strains of *A. baumannii* AB5075 revealed that Aar-mediated repression of *carO* occurred under physiological conditions in this pathogen, and the seed region was again critical for Aar function. Aar-mediated repression of *bfnH* could not be validated in *A. baumannii* using the same experimental setup. It is possible that in the single experimental condition tested, the endogenous levels of Aar were too low for efficient control of the lower affinity target *bfnH* (see Fig. 4C, D). Moreover, while we did not address whether Aar-mediated target control depends on assisting chaperones—a previous study demonstrated a negative influence of Hfq on *carO* mRNA levels in *A. baumannii* ATCC17978 (Kuo *et al*., 2017). Future analysis should therefore address to what extent this effect may be due to Aar and more generally, what role Hfq plays in the post-transcriptional control of gene expression in *Acinetobacter* spp.

What is the biological function of Aar? Aar is expressed in a growth-phase dependent manner, where the highest levels of the sRNA were detected in the stationary phase (**Figure 3C**). Furthermore, Aar is induced in response to several environmental stress conditions, including a temperature shift and osmotic shock (**Supplementary Figure 9**), suggesting a role of Aar in the adaptation of *A. baumannii* to its host environment. Based on our experimental data, Aar fine-tunes the expression of several OMPs, primarily CarO. Generally, bacterial sRNAs that accelerate OMP degradation are often induced by stressful environmental conditions that cause envelope damage or OMP misfolding (Vogel and Papenfort, 2006; Holmqvist and Wagner, 2017; Fröhlich and Gottesman, 2018) and it is plausible that Aar plays a similar role in re-establishing envelope homeostasis in *A. baumannii* during the colonisation of inhospitable environments (Hews *et al*., 2019; Geisinger *et al*., 2019). Further investigations into the regulators of Aar may broaden our understanding of its role in *A. baumannii* and the conditions in which this becomes relevant.

Two primary targets of Aar derived from our study are the mRNAs of the OMP CarO and of the receptor for the baumannoferrin siderophore cluster BfnH. Both CarO and BfnH are central to *Acinetobacter* virulence and pathogenicity therefore Aar might possess a role in *A. baumannii* virulence. Clinical *A. baumannii* strains lacking *carO* displayed increased resistance to carbapenems, leading to the suggestion that CarO acts as a porin for these antibiotics (Limansky *et al*., 2002; Mussi *et al*., 2005), even if this view has recently been challenged (Zahn *et al*., 2015; Geisinger *et al*., 2020). The loss of *carO* in *A. baumannii* ATCC17978 lowered the adhesion to epithelial cells and reduced killing of mice in a sepsis infection model (Labrador-Herrera *et al*., 2020). CarO is also upregulated in *A. baumannii* ATCC17978 biofilms and its disruption impeded biofilm formation (Cabral *et al*., 2011). We discovered that exposing *A. baumannii* to high concentrations of sodium chloride greatly increases Aar expression, as was also reported in *A. baylyi* (Schilling *et al*., 2010). CarO has been established to be the second most abundant protein in the *A. baumannii* OM, and since this porin has cationic affinity, it may lead to excessive uptake of sodium (Siroy *et al*., 2005; Labrador-Herrera *et al*., 2020). The expression of *carO* was reduced at the transcriptional level when *A. baumannii* was cultured in LB with 200 mM of sodium chloride and it was suggested that CarO is shed from the OM into the culture media in the form of outer membrane vesicles when cultured in high concentrations of sodium chloride (Hood *et al*., 2010). Aar may keep CarO levels balanced, potentially reducing *A. baumannii* sensitivity to osmotic stress.

In addition, CarO was shown to be involved in the uptake of L-ornithine and other negatively charged amino acids through passive diffusion (Mussi *et al*., 2007; Zahn *et al*., 2015). Similarly, hydroxymate siderophores like baumannoferrin use L-ornithine as a biosynthetic precursor, suggesting a link between BfnH and CarO (Penwell *et al*., 2015; Sheldon and Skaar, 2020; Cook-Libin *et al*., 2022). The potential involvement of CarO and BfnH in synthesis and uptake of siderophores might imply that Aar contributes to the regulation of iron starvation responses in *A. baumannii*. This is reinforced by the observation that most Aar-*carO* and Aar-*bfnH* chimeric reads were identified in the Hi-GRIL-seq iron starvation condition and by the increase in Aar expression during iron starvation (**Figure 13)** (Schilling *et al*., 2010). Based on the repressive role that Aar exerts on CarO and BfnH, it is thus tempting to speculate that this sRNA plays a broader role in the pathogenicity of *Acinetobacter* species, which could be explored in future studies.

Overall, the work presented here demonstrates that Hi-GRIL-seq is a valuable tool for identification of sRNA-regulated mRNA molecules, particularly in bacterial species where the role of RBPs is unknown or contentious. We present a first global insight into the mechanism of a sRNA-mediated post-transcriptional regulation in *A. baumannii*. Our Hi-GRIL-seq results provide a foundation for further research of *A. baumannii* sRNAs and will facilitate future in-depth functional characterisation.

## DATA AVAILABILITY

The raw sequencing data are available at NCBI under BioProject PRJNA1043178.

## ACKNOWLEDGEMENTS

We are grateful to Siân V. Owen (Harvard Medical School, USA) for providing purified T4 phage lysate and to the labs of Suzana P. Salcedo (University of Wisconsin-Madison, USA), Jay C. D. Hinton (University of Liverpool, UK), Xavier Charpentier and Maria-Halima Laaberki (Inserm, Lyon, France), Paolo Visca (Università Roma Tre, Italy) and Jörg Vogel (Helmholtz Institute for RNA-based Infection Research, Würzburg, Germany) for sharing protocols, plasmids and strains. Deirdre Muldowney (TCD) is acknowledged for technical support. This study has been funded by a Research Grant 2018 from the European Society of Clinical Microbiology and Infectious Diseases (ESCMID) and has emanated from research supported in part by Science Foundation Ireland under Grant number 21/EPSRC/3754 to C. Kröger. A. S. Ershova received funding from the European Union’s Horizon 2020 research and innovation program under the Marie Skłodowska-Curie grant (agreement number 896441). F. Hamrock is supported by a 1252 Postgraduate Research Studentship from Trinity College Dublin, a 2023 Irish Research Council, Government of Ireland Postgraduate Scholarship (GOIPG/2023/4991) and is a recipient of an EMBO Scientific Exchange Grant (no. 9316). A. Shaibah is funded by the Saudi Ministry of Higher Education represented by King Abdulaziz University in Jeddah, Saudi Arabia. M. M. Sulimani is funded by a Custodian of the Two Holy Mosques Scholarship program, Saudi Arabia.

---

**Supplementary Figure 1 – Hi-GRIL-seq in *A. baumannii* AB5075.**

**(A)** Dose-response experiment to identify the concentration of L-arabinose that caused a reduction in cell density without causing excessive cell death was determined. Absorbance of strains carrying the pVRL2Z-*t4rnl* plasmid and strains carrying the empty control vector (pVRL2Z) plasmid were measured following over-night growth (16 h) with different concentrations (from 0 mM to 100 mM) L-arabinose. **(B)** The percentage of RNA species identified by Hi-GRIL-seq in iron starvation (DIP), imipenem shock (IMIP), the induced control (IND) and the uninduced negative control (NI) treatment conditions. Results from independent biological duplicates of each treatment condition is visualised as adjacent bars. The sum of the RNA species percentage is > 100% due to the presence of sequencing reads derived from regions with overlapping annotations.

**Supplementary Figure 2 – Global identification of sRNA-RNA interactions identified by Hi-GRIL-seq in *A. baumannii* AB5075 in different conditions.**

The genomic locations of sRNA-containing chimeric reads ≥ 50 sequencing reads were mapped to the AB5075 chromosome in the (**A**) T4 RNA ligase uninduced (NI) control condition, (**B**) T4 RNA ligase induced (IND) condition, (**C**) iron starvation (DIP) condition and (**D**) imipenem shock (IMIP) condition. The x-axis represents the 5′ end of each chimeric fragment while the y-axis represents the 3′ end of the fragment. Each dot represents the exact genomic coordinate of these fragments. The size of the dot is proportional to the number of sequencing reads observed. The location of all sRNA candidates contained within RNA-RNA chimeras are shown. The locations of the Aar-carO chimeric sequencing reads are labelled in red.

**Supplementary Figure 3 – Chromosomal sRNA-RNA interactions that are overrepresented in the T4 RNA ligase induced conditions (IND, DIP and IMIP).**

Analysis Hi-GRIL-seq revealed that 36 *A. baumannii* AB5075 sRNA candidates were ligated to RNA molecules in other regions of the chromosome. This resulted in the identification of 632 potential chromosomal sRNA-RNA interactions that matched the cut-off scores applied. These interactions are visualised for each sRNA candidate across the AB5075 chromosome.

**Supplementary Figure 4 – Prediction of Aar-mRNA interactions identified by Hi-GRIL-seq identifies a possible Aar seed region.**

The sRNA-mRNA interaction prediction tool IntaRNA was used to identify likely interactions between Aar and Hi-GRIL-seq derived mRNA targets. The position of the IntaRNA predicted interaction for each target is shown on the chromosome for **A**) *ABUW_RS07310* and **B**) *ABUW_RS04325*. The position of the TSSs is shown as curved arrow. The location of the predicted interactions is shown relative to the start codon of mRNA targets. The start codon is in bold and underlined. The predicted Aar “seed” region (5′-CUCC) is highlighted in red.

**Supplementary Figure 5 – Characterising the Hi-GRIL-seq conditions in which *aar*-*mRNA* chimeric reads are most abundant.**

The conditions in which (**A**) *aar*-*carO*, **(B)** *aar*-*bfnH,* **(C)** *aar*-*ABUW_RS07310* and **(D)** *aar*- *ABUW_RS04325* chimeric molecules were most abundant in Hi-GRIL-seq We show the number of chimeras identified in the non-induced (NI) control and the induced (IND), iron starvation (DIP) and imipenem shock (IMIP) conditions. We show the mean of the two biological conditions while the individual points represent the number of chimeras identified in each biological replicate.

**Supplementary Figure 6 – Location of *aar*-*ABUW_RS07310/ABUW_RS04325* chimeric reads mapping to the AB5075 chromosome.**

The location of the **(A)** *ABUW_RS07310* and **(B)** *ABUW_RS04325* portions of Aar-containing chimeric fragments were mapped to the AB5075 for each Hi-GRIL-seq condition. To facilitate visualisation of the chimeric reads, the reads from the two biological replicates of each condition were merged. A coverage profile of the number of chimeric reads is shown on the upper part of each track. Individual chimeric reads are shown in the lower part of each track.

**Supplementary Figure 7 – *aar* forms separate clusters in *A. baumannii* and *A. baylyi*.**

(**A**) Sequence conservation analysis of the *aar* locus in *A. baumannii* and *A. baylyi* strains showing the extent of sequence divergence in *aar*. The location of the predicted *A. baumannii* and *A. baylyi* TSSs and the seed region are shown as arrows. The mutated form of the seed region is indicated. (**B**) Phylogenetic visualisation of *aar* conservation within *A. baumannii* and *A. baylyi*. These organisms form separate groups where a subgroup of *A. baylyi* contains a shorter *aar* sequence.

**Supplementary Figure 8 – Predicting the secondary structure of Aar.**

(**A**) The secondary structure of full-length Aar was predicted using RNAfold and was visualised using VARNA showing the predicted minimum free energy (MFE) structure (Darty *et al*., 2009; Lorenz *et al*., 2011). Individual nucleotides are coloured based on their base-pairing probabilities. The Aar seed region is highlighted. (**B)** The read coverage of the *aar* locus in the Hi-GRIL-seq mapped reads is visualised in JBrowse2 (Diesh *et al*., 2023). The sequencing reads are shown for one biological replicate of each Hi-GRIL-seq condition tested (NI = non-induced control, IND = induced control, DIP = iron starvation and IMIP = imipenem shock). The mapped sequence read data can be viewed at: http://bioinf.gen.tcd.ie/jbrowse2/Hi-GRIL-seq.

**Supplementary Figure 9 – Aar expression is induced by different environmental conditions.** The expression of Aar in *A. baumannii* in several different conditions was compared by Northern blotting. *A. baumannii* was grown to LEP at either 37°C or 25°C. RNA was isolated from cells grown at 37°C that were shocked for 15 min with 0.3 M NaCl (NaCl), 200 µM 2,2-dipyridyl (DIP) or 1 µg/ml polymyxin B (PB) and from a mock treatment control grown at 37°C (37°C). RNA was also isolated from cells following a 15 min temperature shift (25°C to 37°C, TS) and from a mock treatment control grown at 25°C (25°C). Ten µg of total RNA was used for Northern blotting. Expression of Aar and the loading control, 5S rRNA, were detected using their respective DIG-labelled riboprobes. This was performed twice independently, and one representative blot is shown.

**Supplementary Figure 10 – Identification of *aar*-*mRNA* interactions *in vitro* using EMSAs.**

The ability of Aar to interact with **(A)** *ABUW_RS07310* and **(B)** *ABUW_RS04325* and of Aar* to interact with **(C)** *carO*, **(D)** *bfnH*, **(E)** *ABUW_RS07310* and **(F)** *ABUW_RS04325* was assessed using EMSAs. This was accomplished by incubating radiolabelled Aar and Aar* in the absence and in the presence of increasing concentrations (from 8 nM to 1000 nM) of putative mRNA targets.

**Supplementary Figure 11 – Disruption of the Aar seed region inhibits Aar-mediated interactions in vitro.**

EMSAs with 5′-radiolabelled wild-type Aar (WT, black) and seed region mutated Aar (Aar*) with **(A)** *ABUW_RS07310* and **(B)** *ABUW_RS04325* RNA were used to quantify the dissociation constants (*K*_D_) of these targets respectively. The *K*_D_ values were calculated from the mean of two independent replicates. **(C)** The effect of mutating the Aar seed region (Aar*) on base-pairing with *bfnH* and *carO* at the nucleotide level was examined using in-line probing. The regions of nucleotides that were protected in wild-type Aar in the presence of these mRNA interaction partners is highlighted (Regions 1-4). Two ladders were included; a T1 ladder, where the sRNA was cleaved at guanine nucleotides, and a OH ladder, which cleaved the sRNA at the nucleotide level. The Aar seed region nucleotide substitution is visible on the T1 ladder (nucleotides 29, 31 and 32). An uncleaved Aar* control was also included.

**Supplementary Figure 12 – In-line probing reveals the location of nucleotides involved in mRNA base-pairing.**

The use of in-line probing enables determination of the Aar secondary structure in the absence and in the presence of different mRNA interaction partners. **(A)** The secondary structure of wild-type Aar in the absence of mRNA targets revealed that it forms two stem loops. **(B)** Aar is protected from cleavage by *carO* RNA at Region 1 and Region 2. **(C)** Aar is protected from cleavage by *bfnH* RNA at Region 1, Region 2, Region 3 and Region 4, resulting in the unfolding of a stem loop. **(D)** Aar* forms a different secondary structure than wildtype Aar, with a small stem loop encompassing Region 1 (nucleotides 14-17).

**Supplementary Figure 13 – Aar suppresses translation of ABUW_RS07310 and sequesters the *carO* Shine-Dalgarno sequence in a two-plasmid reporter system in *E. coli*.**

**(A)** The involvement of Aar in modulating expression of ABUW_RS07310 was assessed using a two-plasmid system in *E. coli*. The relative fluorescence intensity of strains carrying the pXG10- ABUW_RS07310 plasmid with either the control plasmid (pJV300) or an Aar overexpression vector (pP_L_-Aar) was measured. A strain only carrying pJV300 was also included to measure autofluorescence of *E. coli*. Error bars represent the standard deviation from three independent biological replicates (n=3). Statistical comparisons were performed using One-way ANOVA followed by Tukey’s multiple comparisons test. **(B)** Disruption of *carO* nucleotides complementary (CarO*) to the Aar seed region disrupts fluorescence. The relative fluorescence intensity of strains carrying the pXG10-CarO* plasmid with either the control plasmid (pJV300), an Aar overexpression vector (pP_L_-Aar) or an Aar* overexpression vector (pP_L_-Aar*) was measured. The results from Figure 5A are also shown as comparators. Error bars represent the standard deviation from three independent biological replicates (n=3). Statistical comparisons were performed using One-way ANOVA followed by Tukey’s multiple comparisons test. Differences were considered statistically significant where ns denotes P>0.05, * denotes P<0.05, ** denotes P<0.01, *** denotes P<0.001 and **** denotes P<0.0001.

**Supplementary Figure 14 – Regulation of BfnH by Aar using a translational reporter in *A. baumannii* AB5075.**

The involvement of **(A)** Aar and **(B)** Aar* in modulating translation of BfnH-sfGFP was assessed in *A. baumannii* using a translational reporter. The relative fluorescent intensity of wild-type (WT), Aar- deletion (Δ*aar*) or Aar seed region mutant (*aar*\*) strains of *A. baumannii* AB5075 carrying the pWH1266-BfnH plasmid was compared at LSP. Control strains carrying the pWH1266 plasmid were also measured to quantify the autofluorescence of strains. Error bars represent the standard deviation from six independent biological replicates (n=6). Statistical comparisons were performed using One-way ANOVA followed by Tukey’s multiple comparisons test. Differences were considered statistically significant where ns denotes P>0.05, * denotes P<0.05, ** denotes P<0.01, *** denotes P<0.001 and **** denotes P<0.0001.

