## Supplementary figures and images for "Global analysis of the RNA-RNA interactome in *Acinetobacter baumannii* AB5075 uncovers a small regulatory RNA repressing the virulence-related outer membrane protein CarO"

### Figure S1

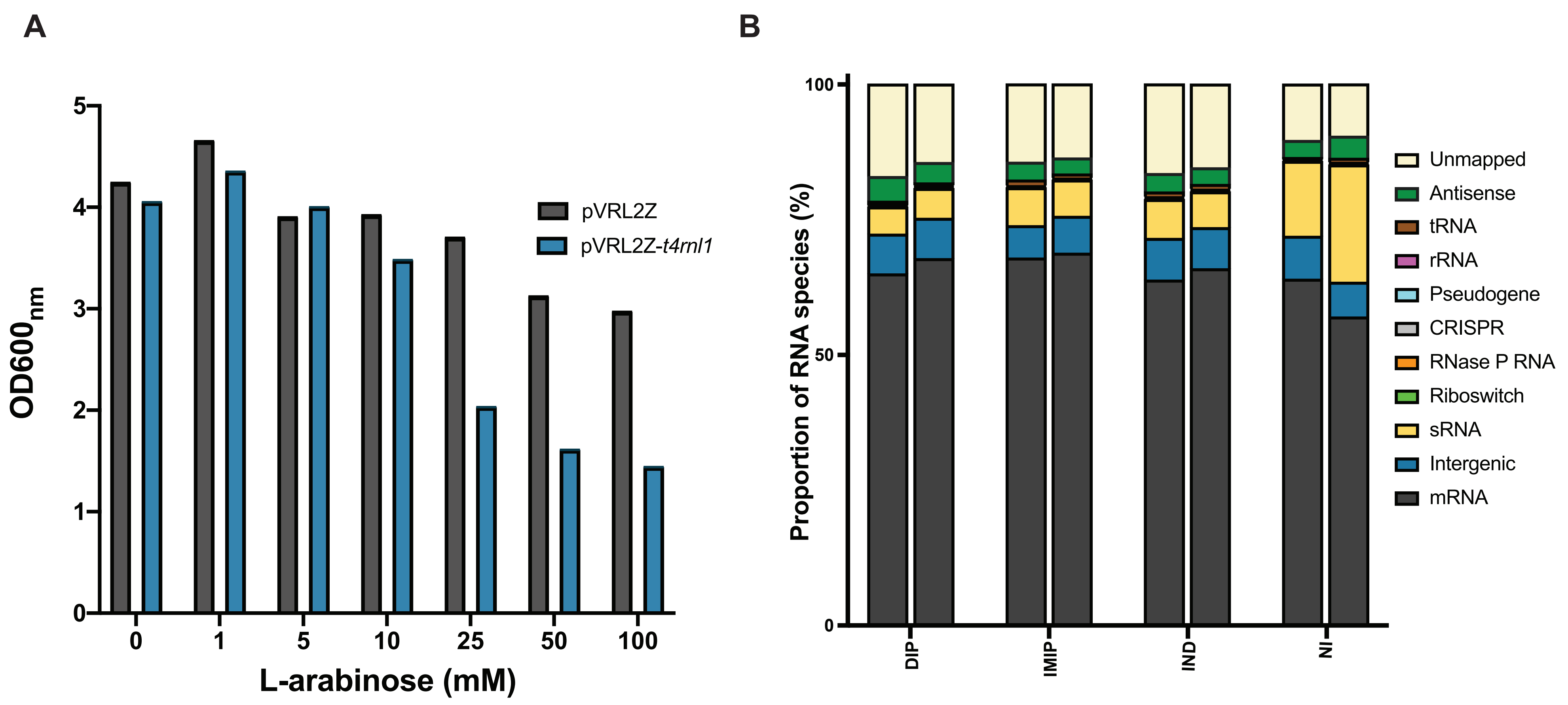

### Figure S2

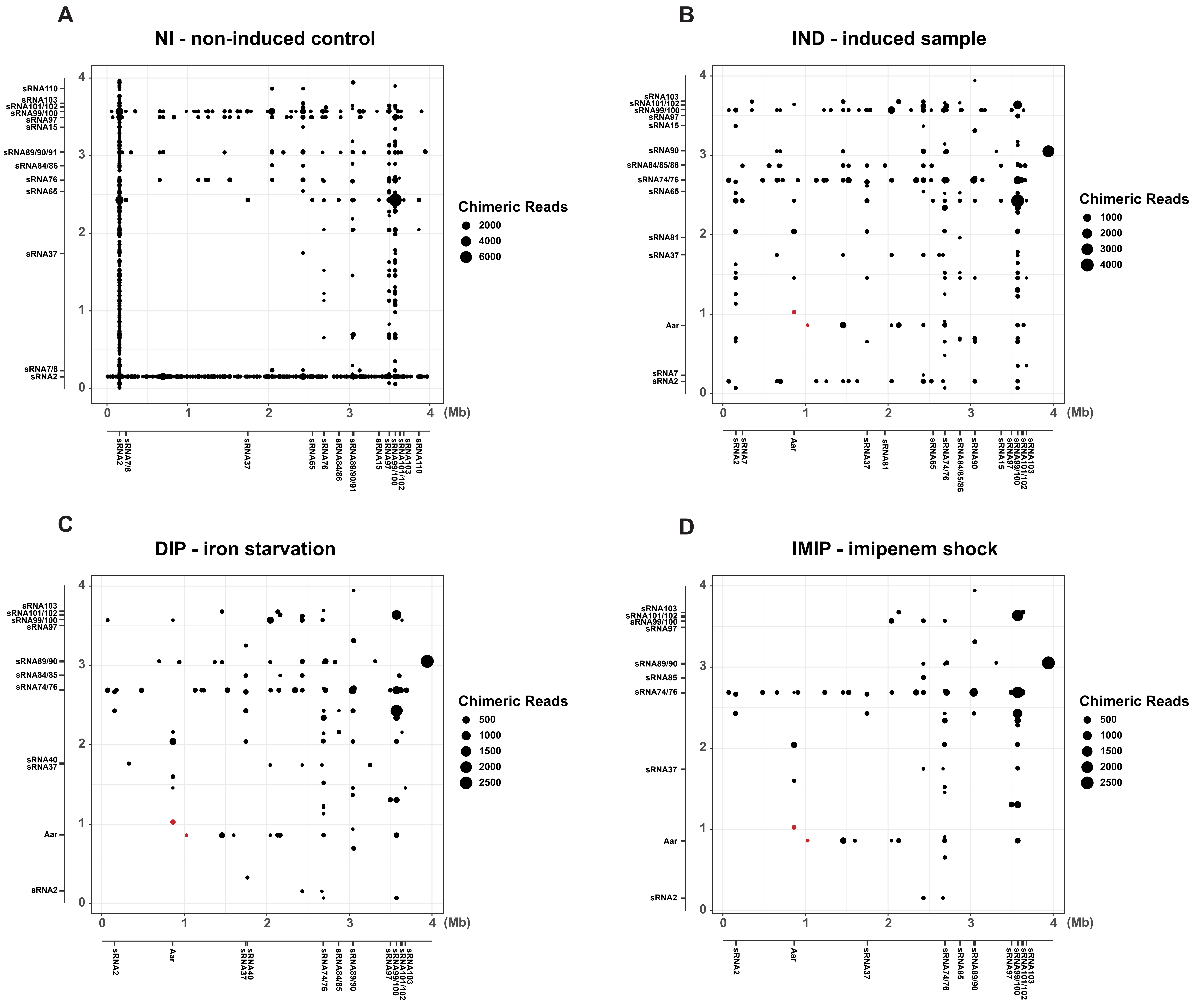

### Figure S3

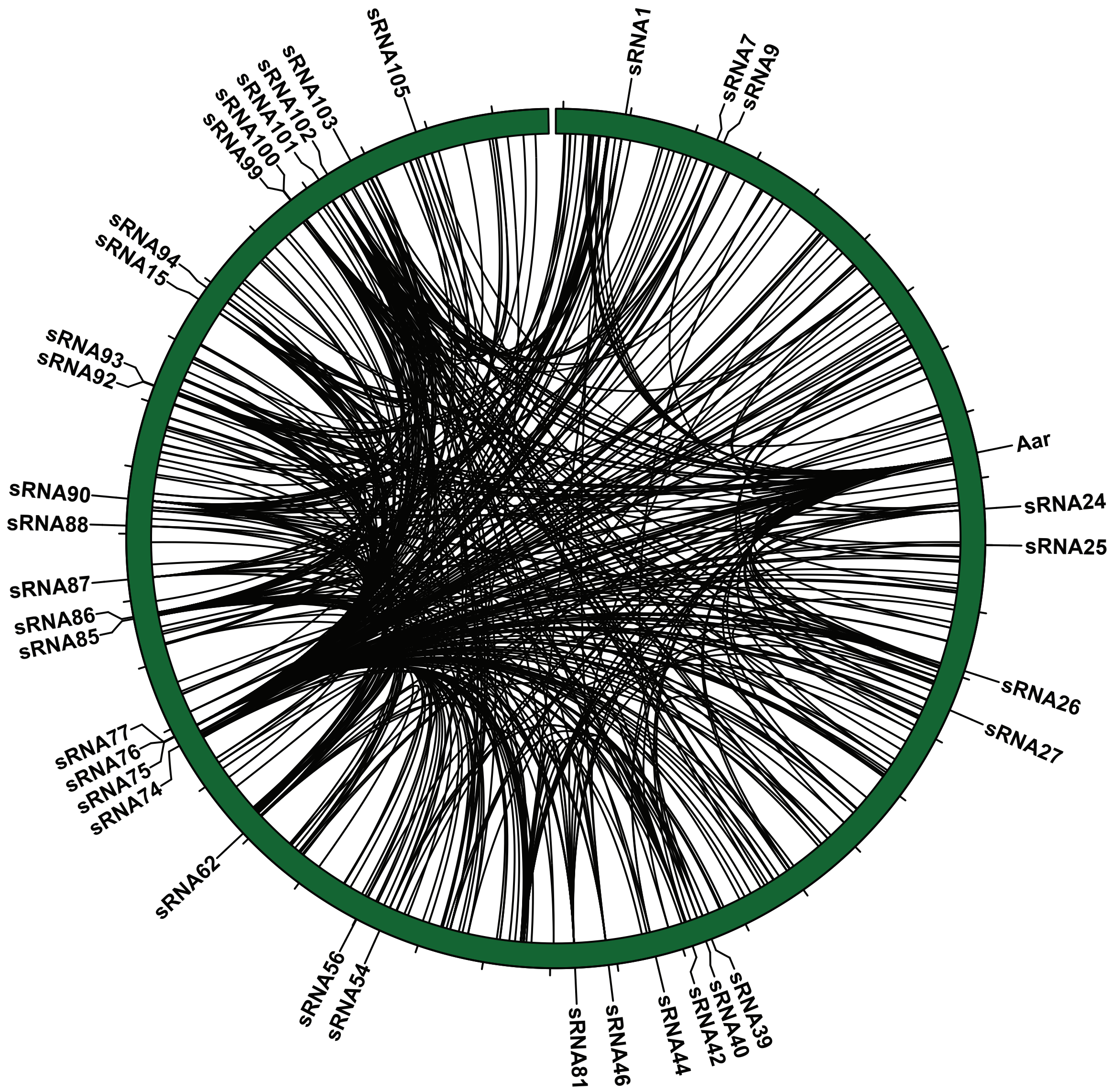

### Figure S4

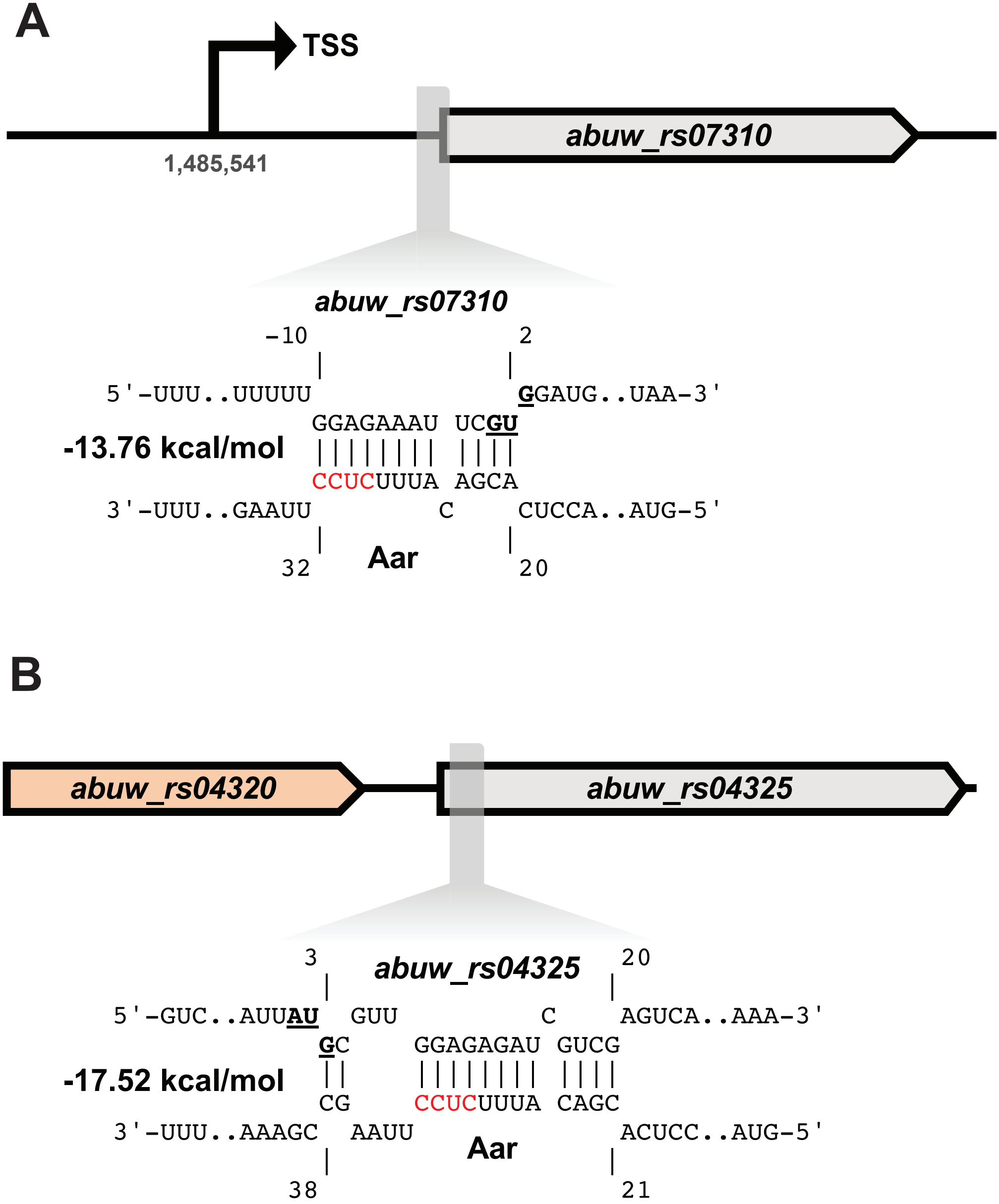

### Figure S5

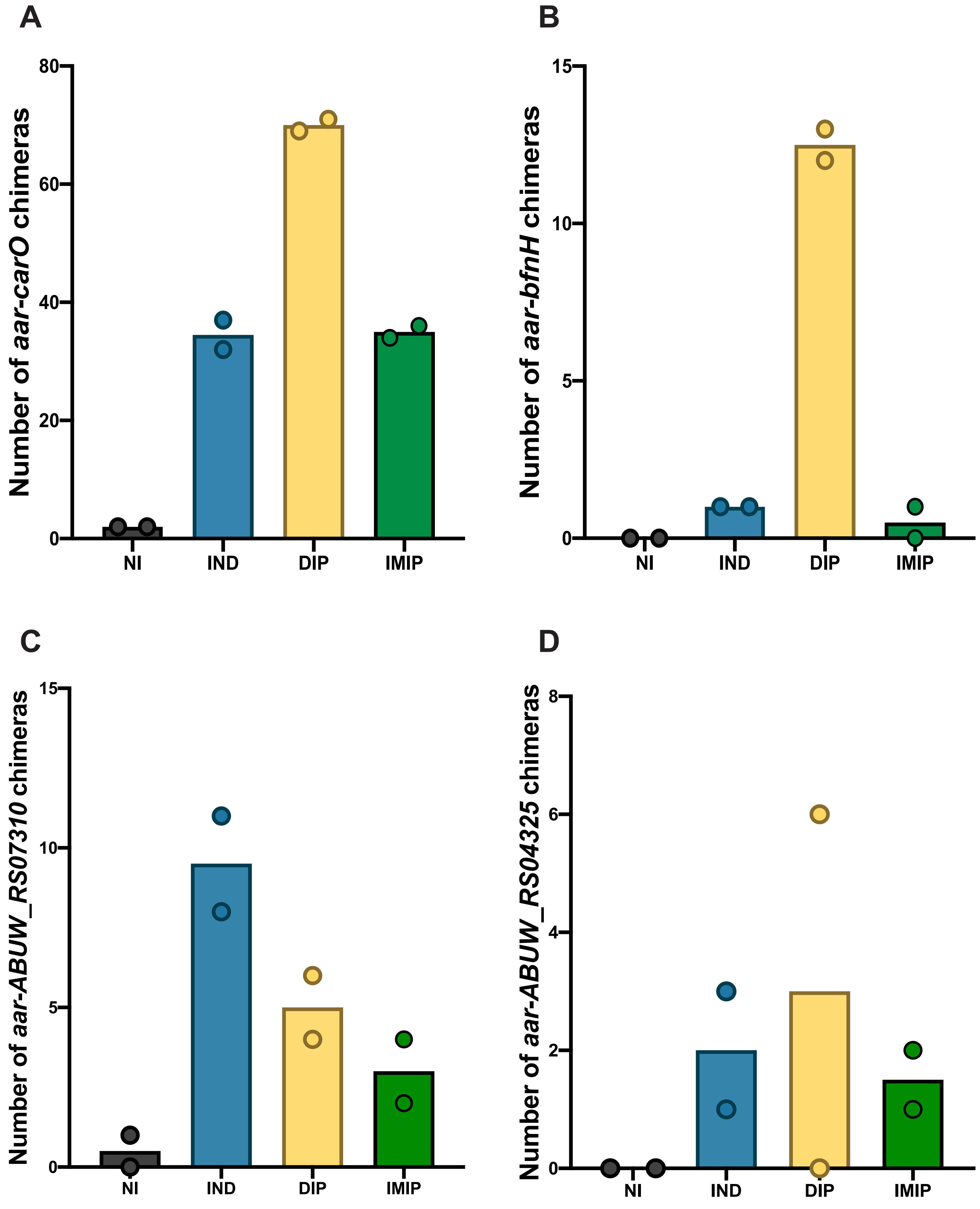

### Figure S6

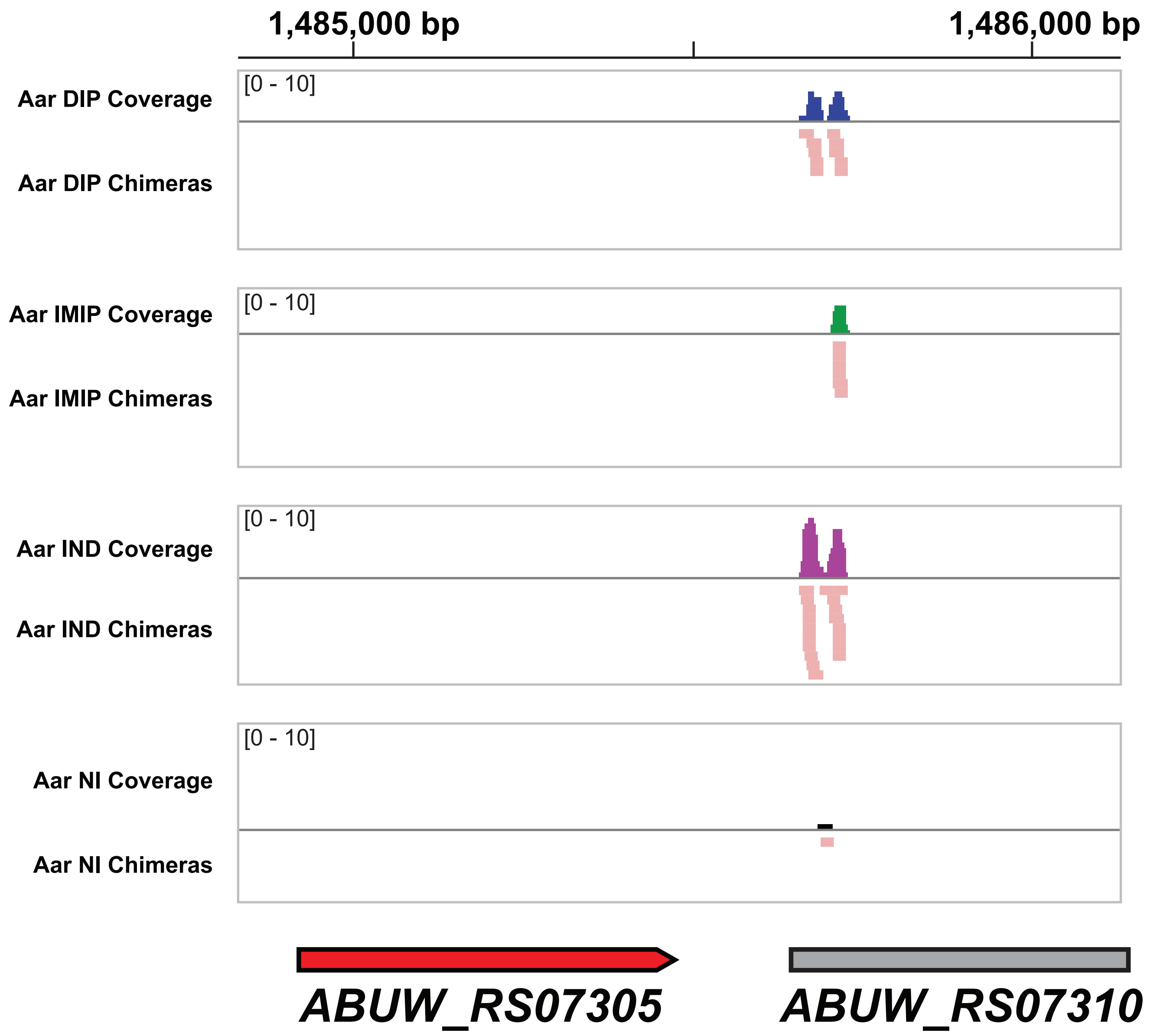

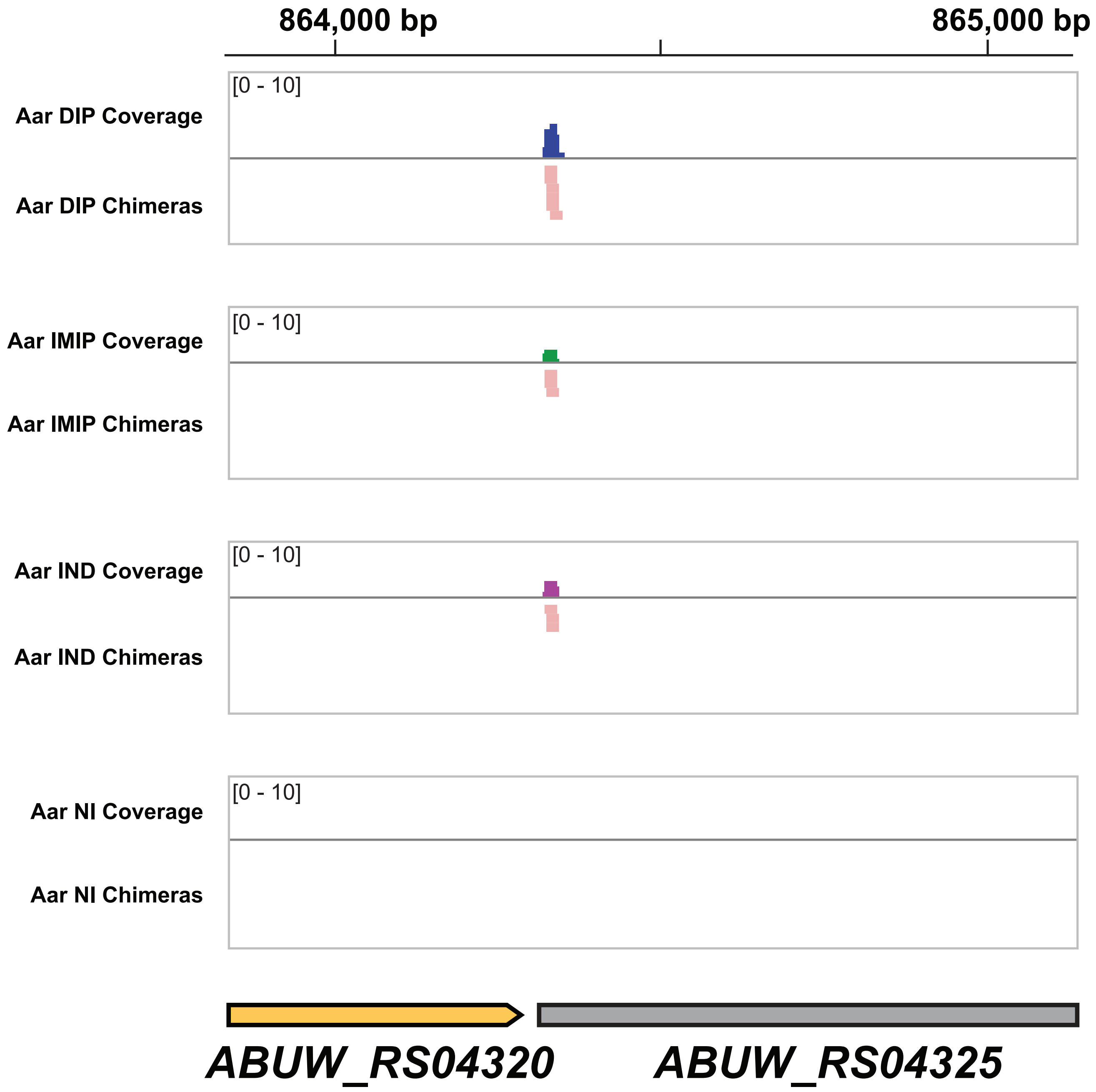

### Figure S7

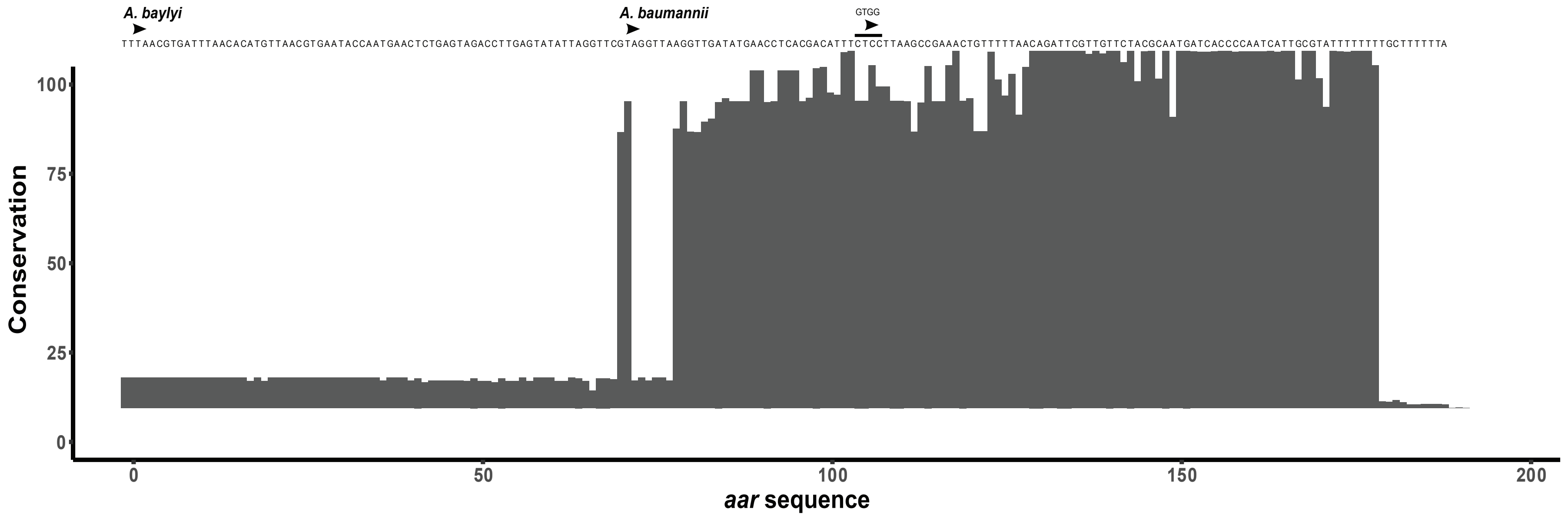

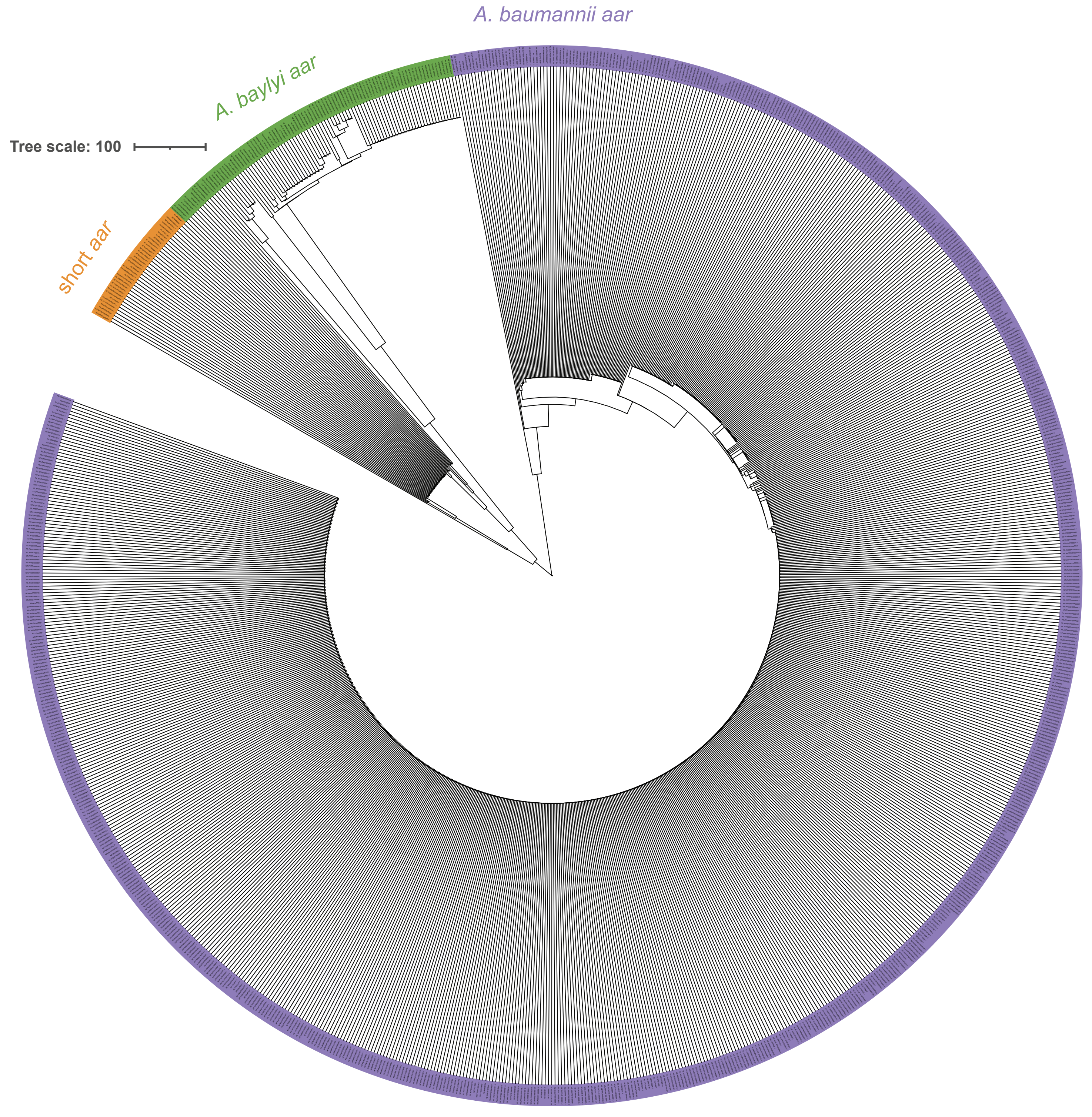

### Figure S8

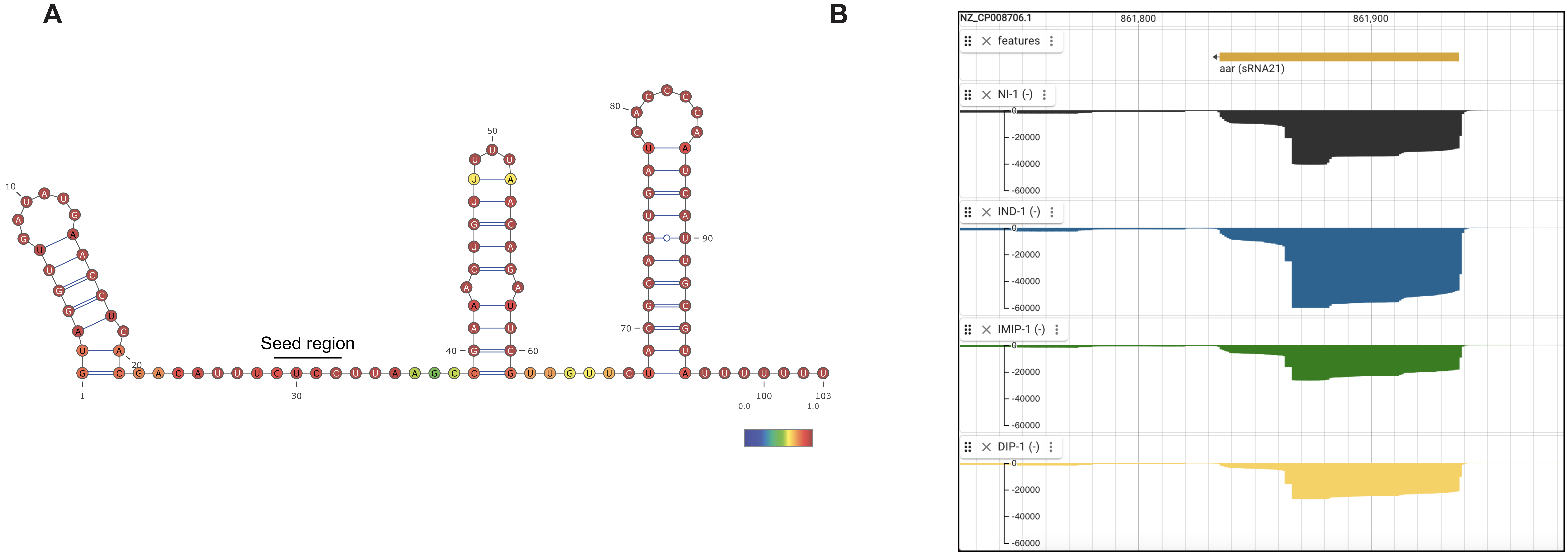

### Figure S9

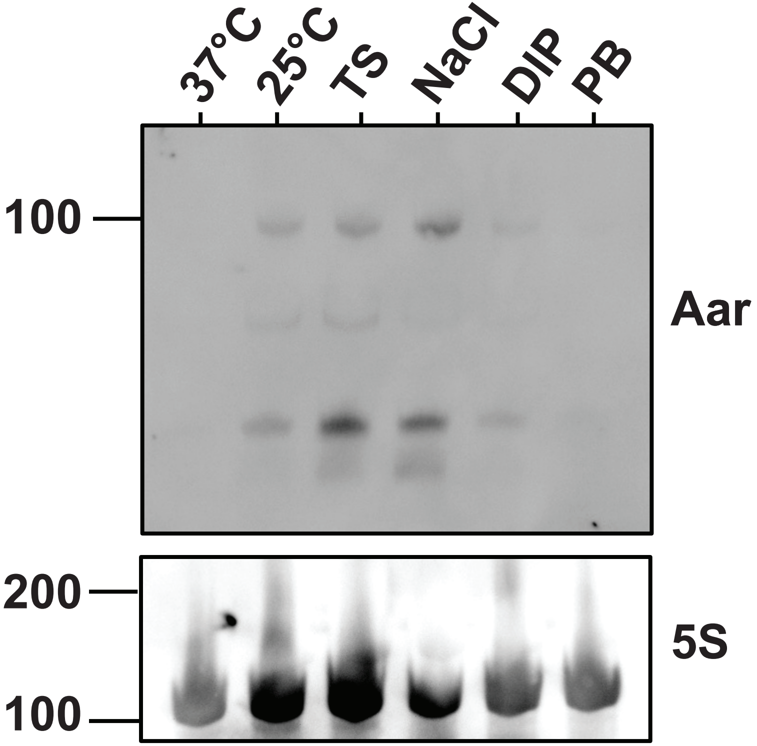

### Figure S10

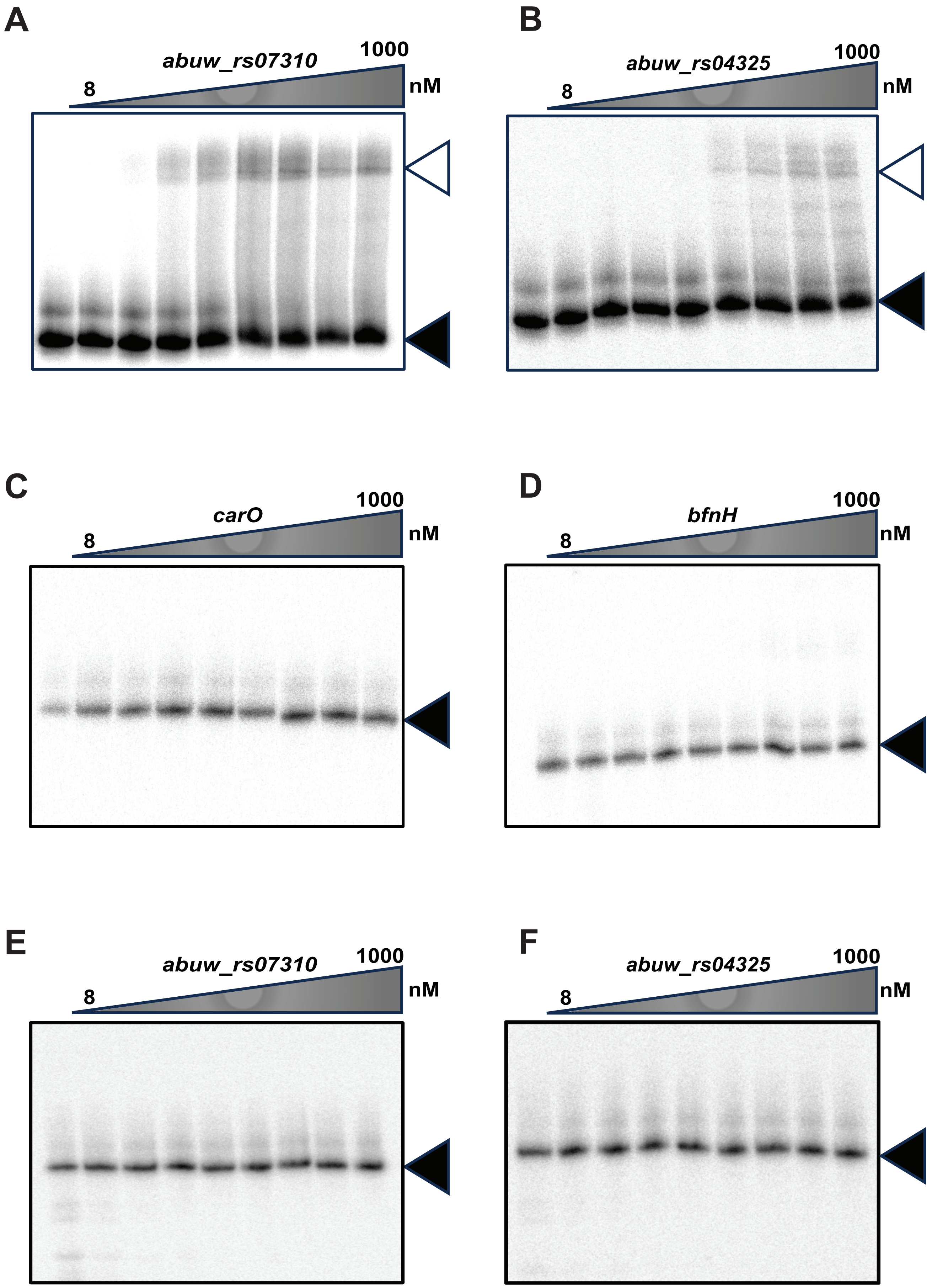

### Figure S11

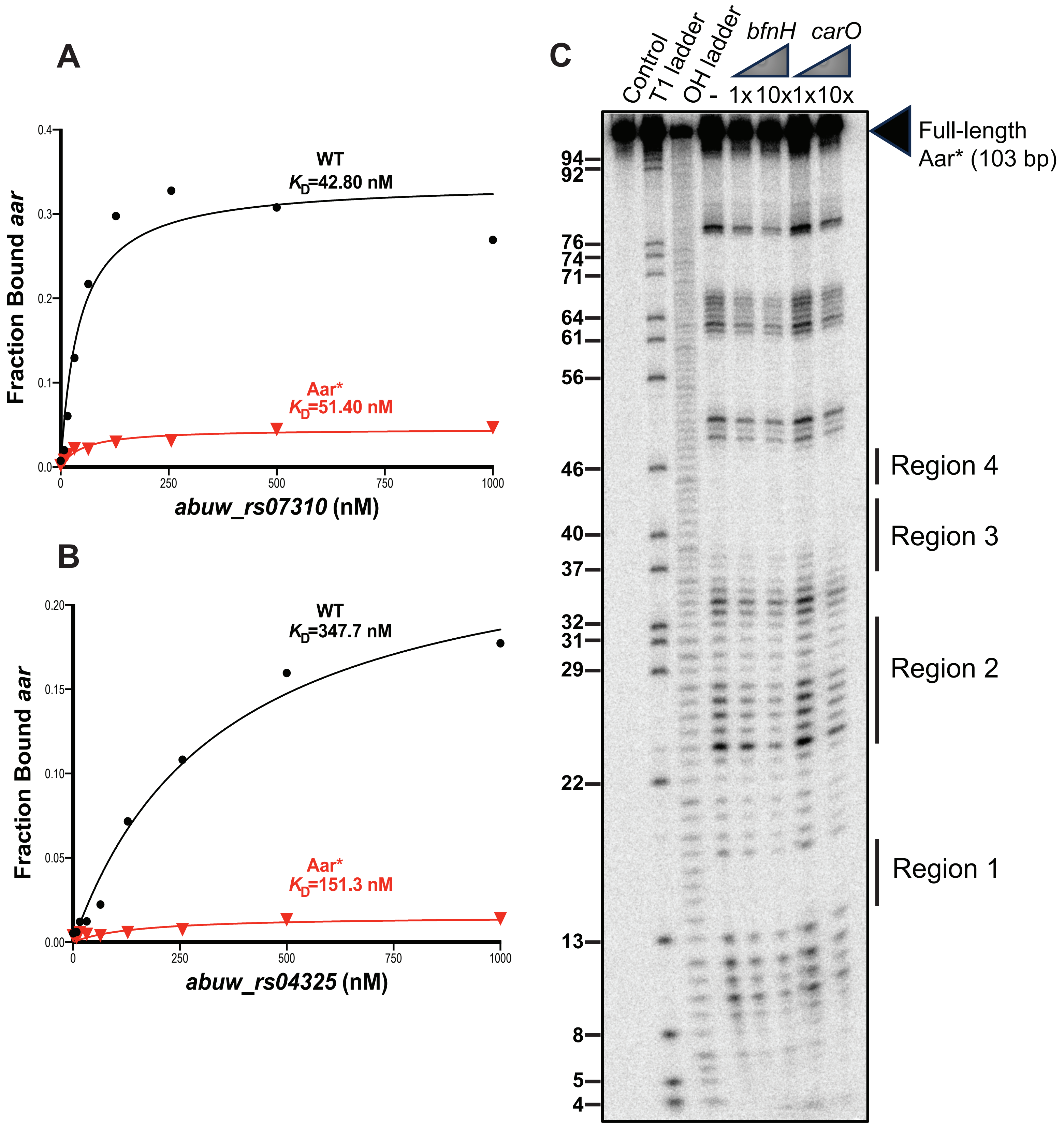

### Figure S12

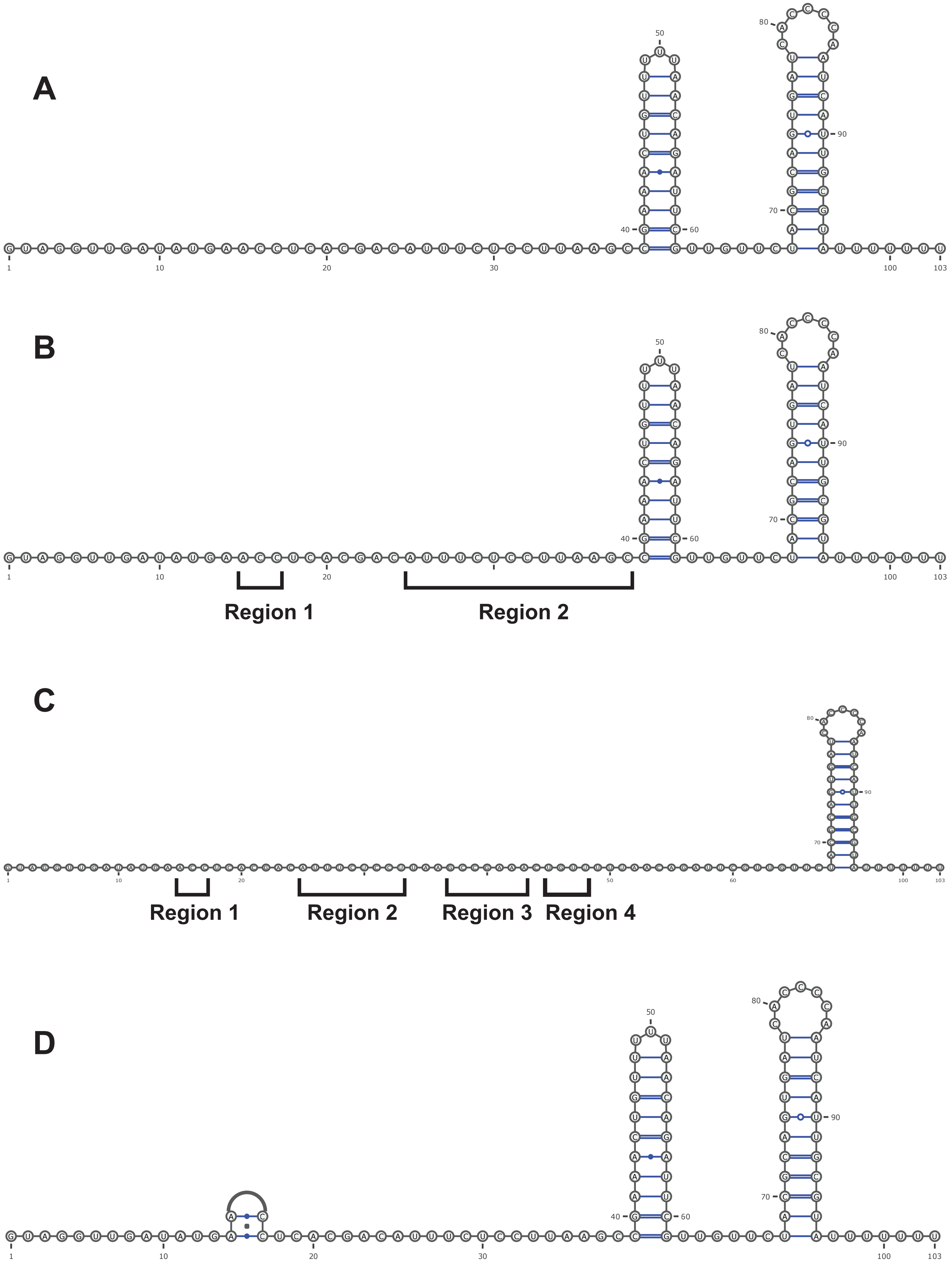

### Figure S13

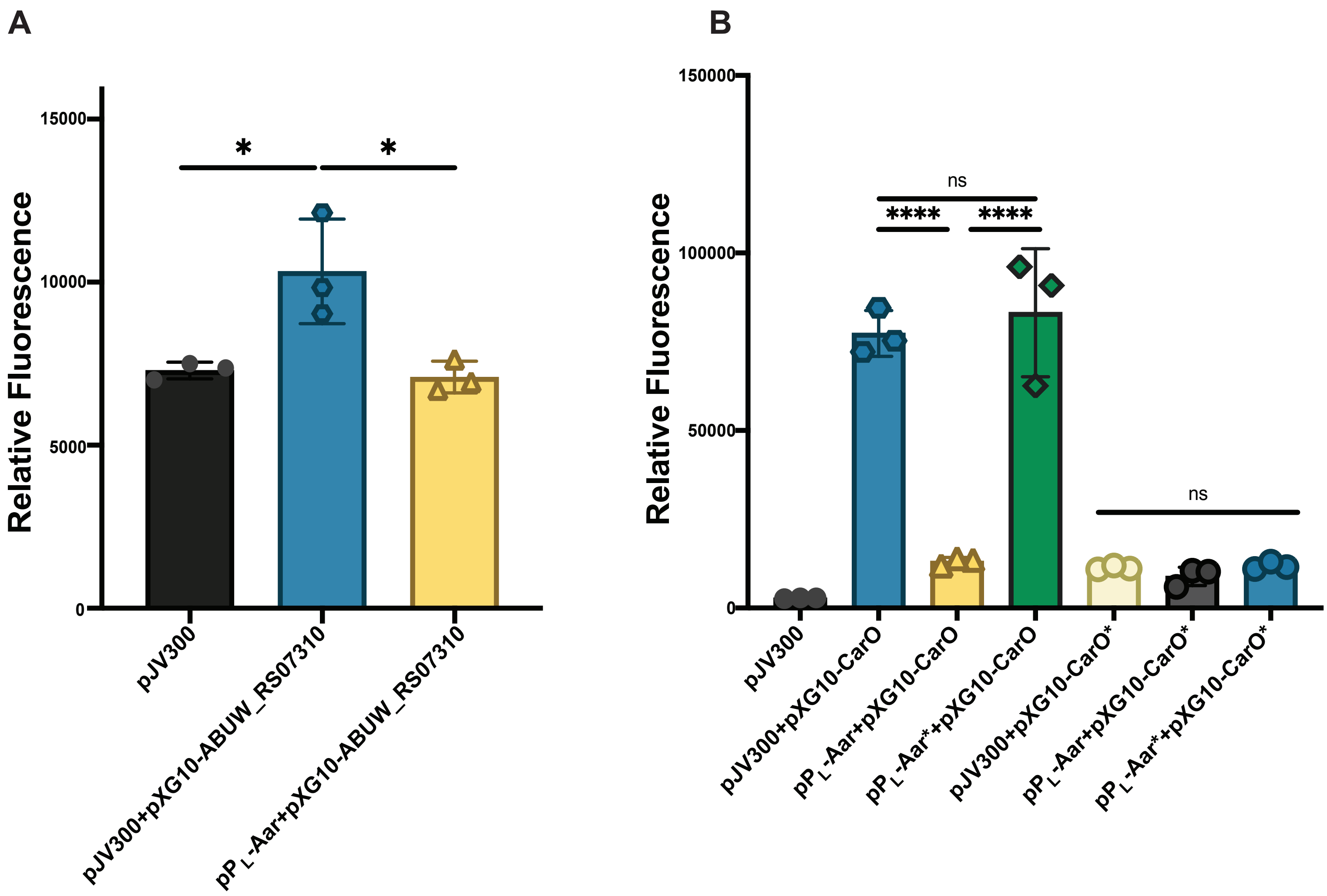

### Figure S14

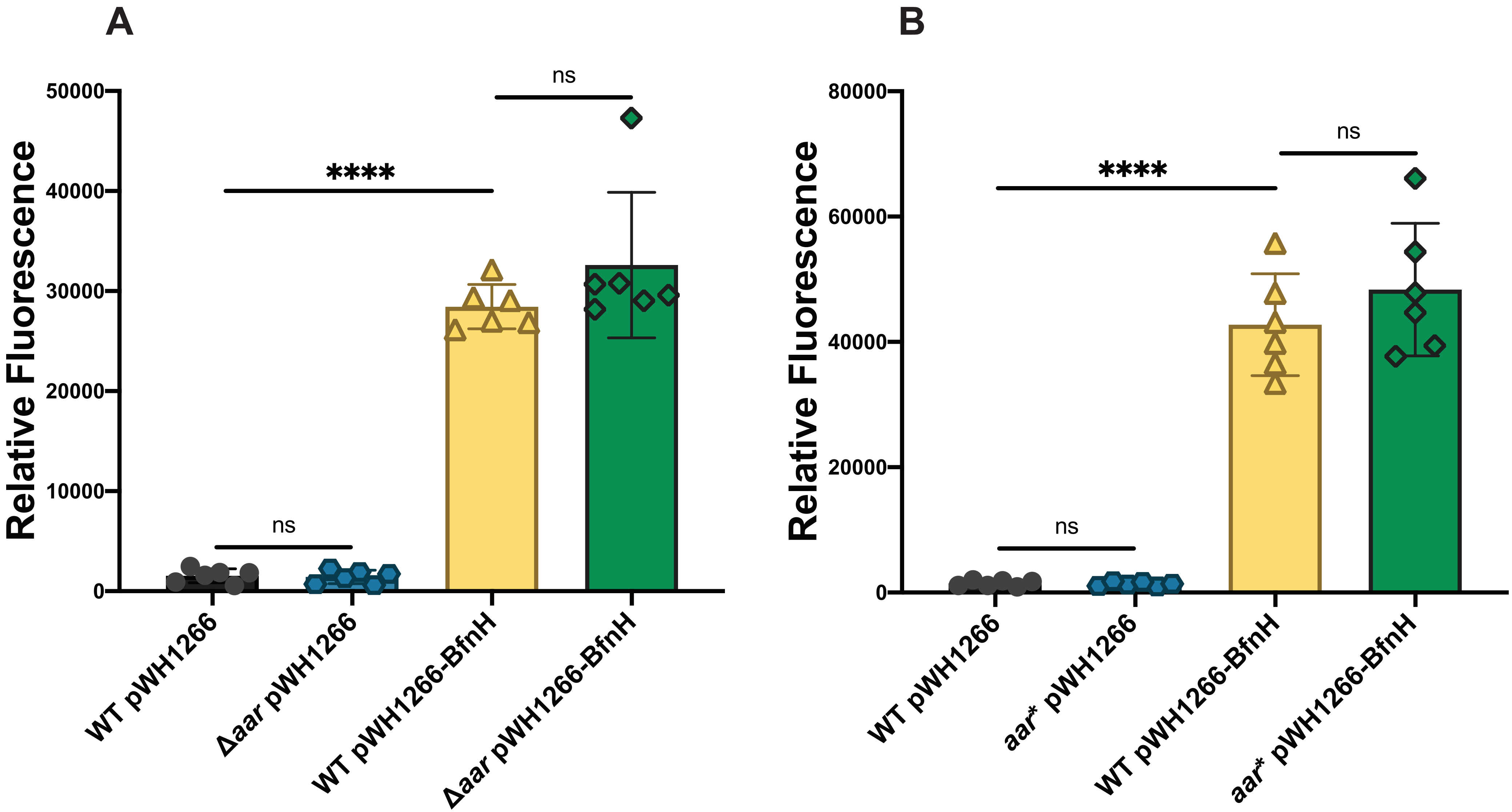
