## Supplementary material for "Global analysis of the RNA-RNA interactome in *Acinetobacter baumannii* AB5075 uncovers a small regulatory RNA repressing the virulence-related outer membrane protein CarO": Table S1

Table S1: Oligonucleotides used in this study. Seed region mutations are underlined.

| **Primer number** | **Name** | **Sequence (5′-3′)** | **Purpose** |
| --- | --- | --- | --- |
| 1 | Aar_P1 | AACTGCAATAATGCTGAGG | Deletion of Aar in *A. baumannii* |
| 2 | Aar_P2 | GGCCCAATTCGCCCTATAGTGAGTCGTGTTCATTAAGCTTCACG | Deletion of Aar in *A. baumannii* |
| 3 | Aar_P3 | GGGTTTGCTCGGGTCGGTGGCATATGTGTCTATAATAATGTTGGC | Deletion of Aar in *A. baumannii* |
| 4 | Aar_P4 | ATTTCTGGTGGTCAAGG | Deletion of Aar in *A. baumannii* |
| 5 | Aar_P5 | CGACTCACTATAGGGCGAATTGGGCC GCTTTCCAGTCGGGAAACCTG | Deletion of Aar in *A. baumannii* |
| 6 | Aar_P6 | CATATGCCACCGACCCGAGCAAACCC CGCCAGGGTTTTCCCAGTCACGAC | Deletion of Aar in *A. baumannii* |
| 7 | Aar_P7 | CGTGAAGCTTAATGAACATGTCTATAATAATGTTGGC | Deletion of Aar in *A. baumannii* |
| 8 | Aar_P8 | TGTTCATTAAGCTTCACG | Deletion of Aar in *A. baumannii* |
| 9 | sRNA21_del_test_F | TGAGTGATAGTGTAACGCG | Testing deletion of Aar in *A. baumannii* |
| 10 | sRNA21_del_test_R | CGAAGTTAAAGATGGTTTAGGC | Testing deletion of Aar in *A. baumannii* |
| 11 | Aar*_overlap_up_R | GGCTTAACCACAAATGTCGTGAGG | Generating Aar seed region mutation (Aar*) |
| 12 | Aar*_overlap_down_F | CACGACATTTGTGGTTAAGCCG | Generating Aar seed region mutation (Aar*) |
| 13 | T4_SalI_F_2 | CGGAGGTCGACATGCAAGAACTTTTTAAC | Amplification of *t4rnlI* gene |
| 14 | T4_HindIII_R_2 | CGGAGAAGCTTTTAGTATCCTTCTGGG | Amplification of *t4rnlI* gene |
| 15 | Aar_pP_L__F | GTGAGCGGATAACAAGATACTGAGCACGTAGGTTGATATGAACC | Cloning Aar into pP_L_ |
| 16 | Aar_pP_L__R | GCCTTTCGTTTTATTTGATGCCTCTAGAAGTTATCAAACAAAGGCGC | Cloning Aar into pP_L_ |
| 17 | BB_pP_L__F | GTGCTCAGTATCTTGTTATCCGCTCAC | Amplification of pP_L_ backbone |
| 18 | BB_pP_L__R | TCTAGAGGCATCAAATAAAACGAAAGGC | Amplification of pP_L_ backbone |
| 19 | Aar*_overlap_up_F | GACGAAGTTAAAGATGGTTTAGGC | Generating Aar seed region mutation (Aar*) |
| 20 | CarO_pXG10_F | ACTGAGCACATGCATATGCTCTTGAATATAATTTTTGTC | Cloning CarO into pXG10sf |
| 21 | CarO_pXG10_R | AGCGGATCCGCTAGCAAGTAAAGCTGTAGTTGTCAC | Cloning CarO into pXG10sf |
| 22 | BfnH_pXG10_F | ACTGAGCACATGCATGCCTCTTGAAAAATCGTTGAG | Cloning BfnH into pXG10sf |
| 23 | BfnH_pXG10_R | AGCGGATCCGCTAGCGTGATTACGTGATTTAACCCC | Cloning BfnH into pXG10sf |
| 24 | ABUW_RS07310_pXG10_F | ACTGAGCACATGCATTTTCTTAATTAAAAATTAACATG | Cloning ABUW_RS07310 into pXG10sf |
| 25 | ABUW_RS07310_pXG10_R | AGCGGATCCGCTAGCTAAATGATGATCATCCACG | Cloning ABUW_RS07310 into pXG10sf |
| 26 | BB_pXG10sf_F | GCTAGCGGATCCGCTGGCTCCGCTGC | Amplification of pXG10sf backbone |
| 27 | BB_pXG10_R | ATGCATGTGCTCAGTATCTCTATCAC | Amplification of pXG10sf backbone |
| 28 | CarO*_overlap_up_F | AAACGTGTTACGTTTAATTGAGG | Generating carO Aar binding site mutation (carO*) |
| 29 | CarO*_overlap_up_R | ATCGTTTTGTGGTTAAGAAAAGGCTCTG | Generating carO Aar binding site mutation (carO*) |
| 30 | CarO*_overlap_down_F | CTTAACCACAAAACGATGAAAGTATTACG | Generating carO Aar binding site mutation (carO*) |
| 31 | CarO*_overlap_down_R | CTGGAATTAATTGGTTTTTATCG | Generating carO Aar binding site mutation (carO*) |
| 32 | pWH1266_CarO_GFP F | AATATTGAAAAAGGAAGAGTATGCTCTTGAATATAATTTTTGTC | Cloning CarO-sfGFP into pWH1266 |
| 33 | pWH1266_BfnH_GFP F | AATATTGAAAAAGGAAGAGTGCCTCTTGAAAAATCGTTGAG | Cloning BfnH-sfGFP into pWH1266 |
| 34 | pWH1266_GFP_R | GAGTAAACTTGGTCTGACAGTTATTTGTAGAGCTCATCCATGC | Cloning target-sfGFP into pWH1266 |
| 35 | pWH1266 B-lactam promoter bb F | ACTCTTCCTTTTTCAATATTATTGAAGC | Amplification of pWH1266 backbone |
| 36 | pWH1266 B-lactam promoter bb F | CTGTCAGACCAAGTTTACTC | Amplification of pWH1266 backbone |
| 37 | Aar_rp_F | GTAGGTTGATATGAACCTCACG | Generating Aar riboprobe |
| 38 | Aar_rp_R | GAATTAATACGACTCACTATAAAAAAAATACGCAATGATTGGG | Generating Aar riboprobe |
| 39 | Aar_in_vitro_F | GTTTTTTTTAATACGACTCACTATAGGGTAGGTTGATATGAACC | *In vitro* transcription of Aar |
| 40 | Aar_in_vitro_R | AAATACGCAATGATTGGG | *In vitro* transcription of Aar |
| 41 | CarO_in_vitro_F | GTTTTTTTTAATACGACTCACTATAGGATGCTCTTGAATATAATTTTTGTC | *In vitro* transcription of carO |
| 42 | CarO_in_vitro_R | CAGCAGCAAGTAAAGCTG | *In vitro* transcription of carO |
| 43 | BfnH_in_vitro_F | GTTTTTTTTAATACGACTCACTATAGGGCCTCTTGAAAAATCGTTGAG | *In vitro* transcription of bfnH |
| 44 | BfnH_in_vitro_R | GTGATTACGTGATTTAACCCC | *In vitro* transcription of bfnH |
| 45 | ABUW_RS07310_in_vitro_F | GTTTTTTTTAATACGACTCACTATAGGTTAGACTGAAAACTCACG | *In vitro* transcription of ABUW_RS07310 |
| 46 | ABUW_RS07310_in_vitro_R | TCCCTGAAAGCAGCATGG | *In vitro* transcription of ABUW_RS07310 |
| 47 | ABUW_RS04325_in_vitro_F | GTTTTTTTTAATACGACTCACTATAGGCTTGATACAAAAAATGTGGCC | *In vitro* transcription of ABUW_RS04325 |
| 48 | ABUW_RS04325_in_vitro_R | AATACTAATACCACCTGCCC | *In vitro* transcription of ABUW_RS04325 |
